# TCF11 Has a Potent Tumor-Repressing Effect than Its Prototypic Nrf1α by Definition of both Similar yet Different Regulatory Profiles, with a Striking Disparity from Nrf2

**DOI:** 10.1101/2021.01.12.426360

**Authors:** Meng Wang, Yonggang Ren, Shaofan Hu, Keli Liu, Lu Qiu, Yiguo Zhang

**Affiliations:** The Laboratory of Cell Biochemistry and Topogenetic Regulation, College of Bioengineering, Chongqing University, No. 174 Shazheng Street, Shapingba District, Chongqing 400044, China; Department of Biochemistry, North Sichuan Medical College, Nanchong 637000, China; School of Life Sciences, Zhengzhou University, No. 100 Kexue Avenue, Zhengzhou 450001, Henan, China

**Keywords:** TCF11, Nrf1α, Nrf2, transcriptomic sequencing, regulatory profiling, hepatocellular carcinoma

## Abstract

Nrf1 and Nrf2, as two principal CNC-bZIP transcription factors, regulate similar but different targets involved in a variety of biological functions for maintaining cell homeostasis and organ integrity. Of note, the unique topobiological behavior of Nrf1 makes its functions more complicated than Nrf2, because it is allowed for alternatively transcribing and selectively splicing to yield multiple isoforms (e.g., TCF11, Nrf1α). In order to gain a better understanding of their similarities and differences in distinct regulatory profiles, all four distinct cell models for stably expressing *TCF11*, *TCF11^ΔN^*, *Nrf1α* or *Nrf2* have been herein established by an Flp-In™ T-REx™-293 system and then identified by transcriptomic sequencing. Further analysis revealed that Nrf1α and TCF11 have similar yet different regulatory profiles, although both contribute basically to positive regulation of their co-targets, which are disparate from those regulated by Nrf2. Such disparity in those gene regulation by Nrf1 and Nrf2 was further corroborated by scrutinizing comprehensive functional annotation of their specific and/or common target genes. Conversely, the mutant TCF11^ΔN^, resulting from a deletion of the N-terminal amino acids 2-156 from TCF11, resembles Nrf2 with the largely consistent structure and function. Interestingly, our further experimental evidence demonstrates that TCF11 acts as a potent tumor-repressor relative to Nrf1α, albeit both isoforms possess a congruous capability to prevent malignant growth of tumor and upregulate those genes critical for improving the survival of patients with hepatocellular carcinoma.

## 1. Introduction

In all life forms, a variety of cell identifications with specialized topological shapes are evolutionarily selectively determined by diverse hub sets of transcription factors (TFs)-regulated gene expression profiles. Thereof, activation or inhibition of distinct TFs is essential for regulation of their target gene expression by binding a specific DNA sequence to maintain and perpetuate the normal and orderly operation of given organisms. In human, there exist over 1600 known and likely putative TFs; they have comprised nearly forty of distinct families [1]. Amongst them, the basic leucine zipper (bZIP) TFs consist of a larger group of the diverse basic-domain superfamily (in accordance with a new classification of TFs as described at http://www.edgar-wingender.de/huTF_classification.html). Of these bZIP factors, the Cap’n’collar (CNC) subfamily members are characterized by a highly conserved 43-aa CNC domain, located N-terminally to the basic DNA-binding domain [2,3]. All CNC-bZIP orthologues share a highly conservatism of evolution from marine bacteria, ascidian, sea urchin, octopus, fly, hydra, worm, bird, insect, fish, frog to mammals including human [4], for an indispensable role in executing the cytoprotective transcriptional responses to changing environmental stress, and thus preserving the cellular homeostasis during development and growth of distinct life forms. Only a functional CNC-bZIP heterodimer with small Maf or other bZIP factors can be allowed for binding to target genes, containing one of specialized antioxidant/electrophile response elements (AREs/EpREs) or other *cis*-regulatory homologous consensus sequences to drive distinct gene expression profiles and shape relevant biological functions.

Notably, Nrf1 [also called NFE2L1 (nuclear factor, erythroid 2 like 1), with its long TCF11 (transcription factor 11) and short Nrf1β/LCR-F1 (locus control region-1) isoforms, Gene ID: 4779] and Nrf2 [also called NFE2L2 (nuclear factor, erythroid 2 like 2), Gene ID: 4780] are two principal CNC-bZIP transcription factors expressed in various cell types and tissues of mammals, including mouse and human [2,5]. Analysis of a neighbor-joining CNC-bZIP phylogenetic tree has unveiled that the membrane-bound Nrf1 orthologues should have emerged at a more ancient stage of the earlier evolution from marine bacteria to humans, and is thereby considered as a living fossil, than the water-soluble Nrf2 [4]. This discovery supports a notion that Nrf1 has a potent capability to fulfill more biological functions far beyond redox regulation that was originally identified. In fact, accumulating evidence reveals that Nrf1 exerts an important role in embryonic development [6,7], osteoblastogenesis [8,9], life quality control of proteostasis by proteasome [10,11] and metabolism [12–14], anti-inflammatory immune response [15], and anti-tumor cytoprotection against hepatoma [16,17], in addition to redox stress defense [18,19]. Conversely, loss of Nrf1’s function by gene-targeting in mice leads to severe oxidative stress and spontaneous development of distinct pathological phenotypes, resembling human non-alcoholic steatohepatitis (NASH) with progressive hepatoma, neurodegenerative diseases or diabetes mellitus [3]. By contrast, Nrf2 is dispensable based on the fact that animal development and growth are unaffected by its functional loss, without any pathological phenotypes. Such being the case, Nrf2 is still accepted as a master regulator of redox homeostasis [20], metabolism [21,22] and DNA repair [23] in order to meet the healthy needs of life. Rather, Nrf2 acts as a two-edged sword to shape significant biological functions in proliferation [24,25], resistance to apoptosis [26,27], angiogenesis [28–30], carcinogenesis [31,32] and invasion [33–35]. Overall, these demonstrate that Nrf1 and Nrf2 elicit similar yet different physiological functions. For instance, our previous work revealed an inter-regulatory crosstalk between Nrf1 and Nrf2 at distinct levels [17], implying that they may regulate each other as a competitive player in similar biological process by distinct ways, but the details require to be further identified.

As a matter of fact, so less attention has been paid on Nrf1 than Nrf2, though the indispensable Nrf1 is highly valued as a robust deterministic transcription factor and identified as an important endoplasmic reticulum (ER) sensor for changes in the intracellular redox, glucose, protein and lipid including cholesterol, status [36,37]. Of note, a unique capability of single *Nrf1* gene confers it to be alternatively transcribed and also further subjected to selective splicing to give rise to multiple isoforms with different tempo-spatial topological properties [38]. Consequently, distinct lengths of Nrf1 isoforms (e.g., TCF11, TCF11^ΔN^, Nrf1α, Nrf1β/LCR-F1) are expressed differentially in distinct type of cells, which makes diverse biological functions of Nrf1 more complicated [3,5,39]. In response to biological cues, the ER-resident Nrf1/TCF11 is topologically dislocated from the lumen into extra-ER compartments, where their glycoproteins are deglycosylated and then subjected to selective juxtamembrane proteolytic processing to yield a mature factor before transactivating target genes (e.g., those encoding proteasomal subunits, antioxidant and cytoprotective proteins). Of note, an isoform longer than Nrf1α was originally referred to as TCF11 [40], consisting of 772 aa, but it is absent in the mouse, while the human prototypic Nrf1α of 742 aa lacks the Neh4L domain, due to alternative splicing of the *TCF11* transcript to remove its exon 4 (Figures 1A, S1A and S1B). As such, Nrf1α retains relative complete structural domains from Neh1L to Neh6L, of which equivalents exist in Nrf2 [3,43]. Thereby, it is postulated that Nrf1α and TCF11 should be two main players to exert differential transcriptional regulation of distinct Nrf1-target genes, but this remains to be proved. In addition, the short isoform Nrf1β [41,42], which was early designated as LCR-F1, lacks the N-terminal domain (NTD) and its adjacent acidic domain 1 (AD1), relative to Nrf1α or TCF11 (Figure S1A).

**Figure 1.**
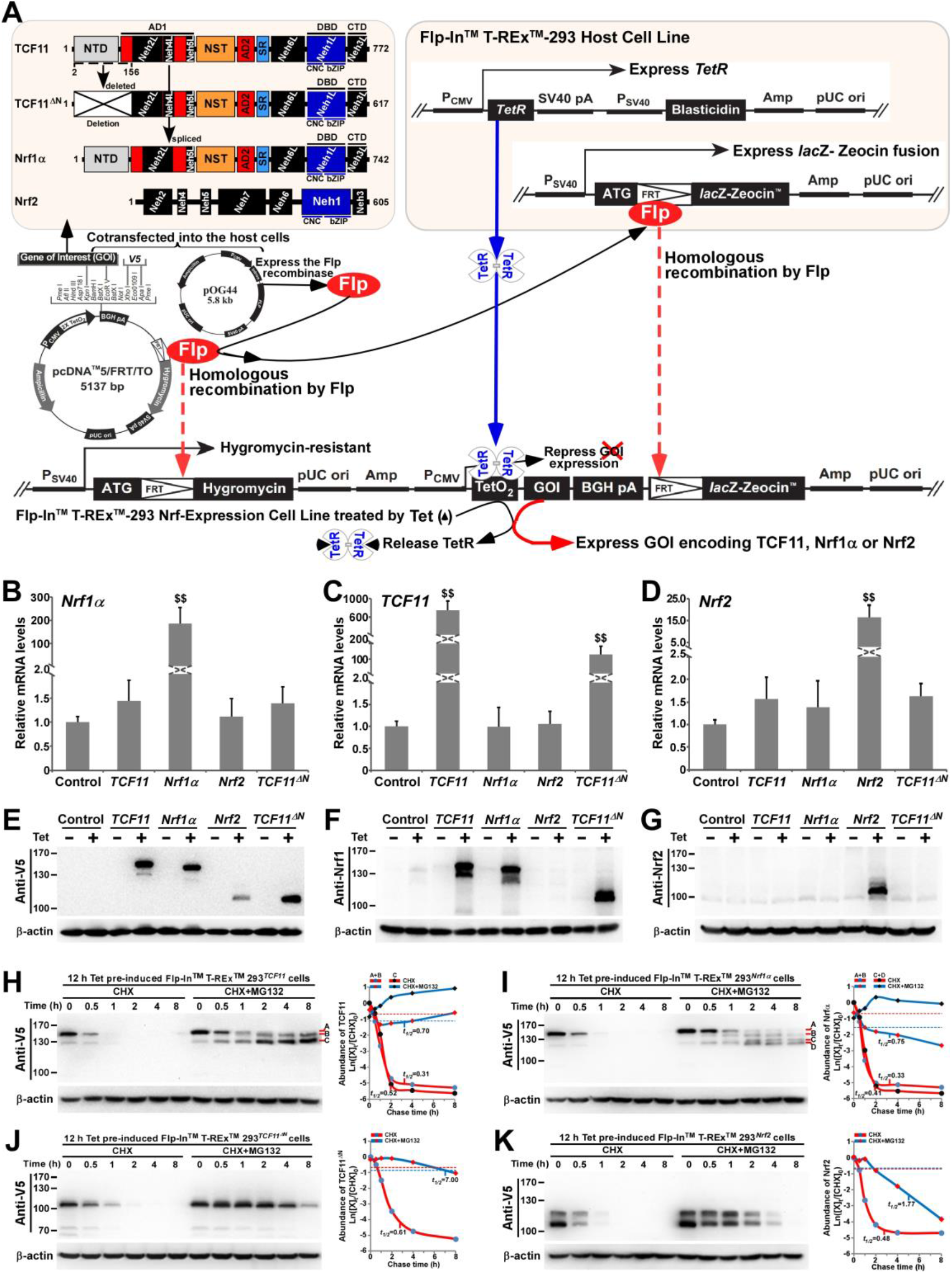
Establishment of four distinct model cell lines to stabilize expression of *TCF11*, *TCF11^ΔN^, Nrf1α* and *Nrf2*. (**A**) A schematic diagram of the Flp-In™ T-REx™-293 system. The system allows the Flp recombinase-mediated homologous recombination of each indicated pcDNA5/FRT/TO-V5 expression constructs (for TCF11, TCF11^ΔN^, Nrf1α or Nrf2) with the Flp-In™ T-REx™-293 host cells through the FRT sites. (**B-D**) After incubation of *TCF11*, *Nrf1α*, *Nrf2* or *TCF11^ΔN^*, as well as control cell lines with 1 μg/mL Tet for 12 h, total RNAs were isolated and then reversely transcribed into the first strand of cDNA. Subsequently, quantitative real-time PCR was employed to identify the mRNA expression levels of *Nrf1α* (B), *TCF11* (C), *TCF11^ΔN^* (C) and *Nrf2* (D) in each of indicated cell lines. The data are shown as mean ± SEM (n = 3 × 3, $$, *p* < 0.01, when compared to the *Control*). (**E-G**) Total lysates of each cell line that had been treated with 1 μg/mL Tet (+) or not (−) were subjected to protein separation by SDS-PAGE gels, and then visualized by immunoblotting with distinct primary antibodies against V5, Nrf1 or Nrf2 to identify the protein levels of TCF11, Nrf1α, Nrf2 and TCF11^ΔN^. (**H-K**) Total lysates of experimental cells, which had been induced with 1 μg/mL Tet for 12 h before being treated with CHX (50 μg/mL) alone or in combination with MG132 (10 μmol/L) for distinct times as indicated, were resolved by SDS-PAGE and then analyzed by Wester blotting with V5 antibody to identify the stability of TCF11 (H), Nrf1α (I), TCF11^ΔN^ (J) and Nrf2 (K) respectively.

To date, growing evidence indicates that Nrf1 and Nrf2, as two versatile leading players in maintaining cellular homeostasis, are essential for important pathophysiological processes in human diseases. Yet, which specific target genes are regulated by both CNC-bZIP factors, and specific biological processes in which such genes are implicated, require for further in-depth study, albeit two recent reports also revealed different portions of between the indicated NRF-target expression profiles [44,45]. Herein, to refine distinct functions of Nrf1, it remains important to distinguish TCF11 and its N-terminally-truncated TCF11^ΔN^ (which is derived from deletion of amino acids at the 2^nd^ to 156^th^ positions in TCF11, and can also occur naturally with the reminiscent Nrf2-like structural domains) from Nrf1α and Nrf2. As for this end, we have established four different cell lines stably expressing *TCF11, TCF11^ΔN^, Nrf1α* or *Nrf2*, respectively, by using an Flp-In™ T-REx™-293 system (as deciphered in Figures 1A & S2). When required, this controllable system is turned on by tetracycline to induce each factor-specific transcriptional expression, while all others are unaffected in each of the indicated cell lines. Subsequently, transcriptomic sequencing of these cell modes unraveled that those common genes regulated by Nrf1 and Nrf2 are more responsible for regulating transcriptional expression and signal transduction in response to stimulus and diseases. The differentially expressed genes regulated by Nrf1, but not Nrf2, are preferentially enriched in carbohydrate metabolism and cellular processes, while Nrf2-specific genes strikingly prefer to developmental process. Notably, Nrf1α and TCF11 share similar regulatory patterns, but they are disparate from those of Nrf2 or TCF11^ΔN^. Besides, TCF11 can also mediate more target genes that are different from those regulated by Nrf1α, displaying distinct biological functions in development and regeneration. This notion is evidenced by our further supportive experiments, demonstrating that TCF11 can serve as a potent tumor-repressor relative to prototypic Nrf1α, albeit both factors possess a congruous capability to prevent tumor growth, as accompanied by up-regulation of those target genes significantly improving the survival rate of patients with hepatocellular carcinoma (HCC).

## 2. Materials and Methods

### 2.1. Chemicals and Antibodies

All chemicals were of the highest quality commercially available. Hygromycin B and blasticidin were purchased from Invitrogen Ltd, which were employed as double drug-screening to select putative positive clones from those transfected Flp-In™ T-REx™-293 expression cells. Tetracycline from Sangon Biotech Co (Shanghai, China) was utilized as an inducible reagent at a concentration of 1 μg/mL. Both cycloheximide (CHX) and MG132 were purchased from Sigma-Aldrich (St Louis, MO, USA). The antibody against Nrf1 proteins was acquired from our own lab (as indicated in Zhang’s [46]), and all other antibodies were also employed against a V5 epitope (Invitrogen), Nrf2 (Abcam), CHP2 (Sangon Biotech), CPS1 (Abcam), FOXO1 (Cell Signaling Technology), IRS4 (Abcam), NKX2-8 (Sangon Biotech), AKR1B10 (Abcam), EPO (Proteintech Group), MUTYH (Sangon Biotech), PKM (Sangon Biotech), GP73 (Proteintech Group), GPC3 (Proteintech Group), Histone H3 (Bioss) or α-Tubulin (Beyotime), whilst β-actin and secondary antibodies were from ZSGB-BIO (Beijing, China).

### 2.2. Cell lines, Cell culture and Transfection

These cell lines expressing *TCF11*, *TCF11^ΔN^*, *Nrf1α*, *Nrf2*, as well as an empty control, were established by using the Flp-In™ T-REx™-293 system (Invitrogen) (Figure S2). Their cDNA fragments, encoding human TCF11, TCF11^ΔN^ (with a deletion to remove the 2^nd^ to156^th^ residues prior to the Neh2L subdomain from TCF11), Nrf1α and Nrf2, respectively, were cloned into pcDNA5/FRT/TO-V5 expression vector, before being cotransfected with a Flp-expressing pOG44 plasmid into the Flp-In™ T-REx™-293 host cells. The Flp recombinase was allowed for its homologous recombination at FRT (Flp Recombination Target) sites existing in the host cells with each of the expression vectors. Of note, an empty expression vector was cotransfected into the host cell, to generate a negative control cell line. Then, the positive expression clones, as indicated, were selected by co-treatment of 150 μg/mL hygromycin B and 15 μg/mL blasticidin. All positively-selected cell lines were allowed for stably expression of target genes, beyond the negative control line, and thus referred to simply as *TCF11*, *TCF11^ΔN^*, *Nrf1α*, *Nrf2* or *Control*, respectively. Moreover, HepG2 was obtained originally from ATCC (Zhong Yuan Ltd., Beijing, China); MHCC97H and MHCC97L were obtained originally from the Live Cancer Institute, Fudan University of China; HL-7702, SMMC-7721 and QGY-7701 were obtained originally from National Infrastructure of Cell Line Resource (NICR), and Huh7 was obtained originally from Japanese Collection of Research Bioresources (JCRB), while both *Nrf1α^−/−^* and *Nrf2^−/−^* cell lines were created from wild-type HepG2 cells [16,17]. Notably, the fidelity of these cell lines had been conformed to be true by their authentication profiling and STR (short tandem repeat) typing maps (which were carried out by Shanghai Biowing Applied Biotechnology Co., Ltd, Shanghai, China) [37].

All experimental cells, except elsewhere indicated, were allowed for growth in DMEM basic medium (GIBCO, Life technologies), with being supplemented with 10% (v/v) foetal bovine serum (FBS, Biological Industries, Israel) and 100 units/mL of either of penicillin and streptomycin, in the 37℃ incubator with 5% CO_2_. Of note, those expression constructs for human Nrf1α, TCF11 and Nrf2 were made by inserting each of their cDNA-encoding sequences into a pcDNA3 vector, respectively. The primer pairs used for this study were provided as shown in Table S1. The cell transfection with one of those indicated plasmids, alone or in combination, were carried out by using Lipofectamine^®^3000 Transfection Kit (Invitrogen) for 8 h, and then allowed for a 24-h recovery from transfection in the fresh medium before being subjected to the indicated experiments.

### 2.3. Quantitative Real-Time PCR

Each of experimental cell lines was subject to its total RNAs isolated by employing an RNAsimple Kit (Tiangen Biotech CO. LTD, Beijing, China). Total RNAs (1 μg) were added in a reverse-transcriptase reaction to yield the first strand of cDNAs (by another RevertAid First Strand Synthesis Kit, from Thermo), which served as the template of quantitative PCR in the GoTaq^®^ qPCR Master Mix (Promega). Then, each pairs of all forward and reverse primers (as listed in Table S1) were also added in an indicated PCR, that was carried out in the following conditions at 95°Cfor 3 min, followed by 40 cycles of 15 s at 95°C, and the last 30 s at 60°C. The final melting curve was validated to examine the amplification quality, and β-actin at its mRNA expression levels served as an internal control for normalization.

### 2.4. Western Blotting

Each of experimental cell lines was harvested in a lysis buffer (0.5% SDS, 0.04 mol/L DTT, pH 7.5) containing the protease inhibitor EASYpacks (Roche, Germany). The lysates were denatured immediately at 100°C for 10 min, sonicated sufficiently, and diluted in 3 × loading buffer (187.5 nmol/L Tris-HCl, pH 6.8, 6% SDS, 30% Glycerol, 150 nmol/L DTT, 0.3% Bromophenol blue) at 100°C for 5 min. Subsequently, equal amounts of protein extracts were subjected to separation by SDS-PAGE containing 4-15% polyacrylamide, followed by immunoblotting with each of distinct antibodies as indicated. On some occasions, the blotted membranes were also stripped for 30 min and then re-probed with an additional antibody, whilst β-actin served as an internal control to verify equal loading of protein in each of electrophoretic wells.

### 2.5. Transcriptome Sequencing Analysis

Total RNAs extracted from each of cell lines, that had incubated with 1 μg/mL of tetracycline for 12 h, were subjected to the transcriptome sequencing by Beijing Genomics Institute (BGI, Shenzhen, China) on an Illumina HiSeq 2000 sequencing system (Illumina, San Diego, CA). All detected mRNAs were fragmented into short fragments (~200bp). The clean reads were obtained during data filtering to remove the low-quality reads, and subjected to sequence mapping to the reference of human genome (GRCh37/hg19 from UCSC database) by using SOAP2 [47]. The resulting expression levels of given genes were calculated by the RPKM method [48]. All those differentially expressed genes (DEGs) were identified, with the criteria fold changes ≥ 2 or ≤ 0.5 and FDR (false discovery rate) ≤ 0.001, by the Poisson distribution model method (PossionDis) [49,50]. Such sequencing metadata have also been submitted to NCBI SRA (PRJNA501789). In addition, the DEGs were functional annotated by using the online tool DAVID (https://david.ncifcrf.gov/) to search their involved GO (gene ontology) terms and pathways, which were further classified with QuickGO (https://www.ebi.ac.uk/QuickGO/) and KEGG (https://www.kegg.jp/) databases.

### 2.6. Lentivirus-Mediated Restoration of Nrf1α or TCF11

One of the lentiviral-mediated expression constructs for *Nrf1α* or *TCF11*, that was designed by a help with supplier (GeneCopoeia, Guangzhou, China), together with the GFP-expressing lentiviral control vector, were co-transfected into our *Nrf1α^−/−^* cells, to establish *Nrf1α*-restored and *TCF11*-restored cell lines. Briefly, the lentiviral-packaging 293T cells (1 × 10^6^) were seeded in a 10-cm dish and cultured in 10 mL DMEM supplemented with 10% FBS. Then, the mixture of 2.5 μg of each lentiviral ORF expression plasmid and 0.25 μg of the Lenti-Pac HIV plasmid in 15 μL of EndoFectin Lenti was incubated with 200 μL of Opti-MEM^®^ (Invitrogen). The DNA-EndoFectin Lenti complex was directly added into cultured cells before being allowed for overnight incubation at 37°C in a CO_2_ incubator, followed by replaced by fresh medium supplemented with 5% FBS. Subsequently, a 1:500 volume of the TiterBoost reagent was further added to the above cultured media and allowed for continuous culture. The pseudovirus-containing culture media for 48 h post-transfection were collected by centrifuging as 500 × *g* for 10 min. Then, the resulting lentivirus titer was estimated, prior to being subjected to efficient transfection of *Nrf1α^−/−^* hepatoma cells.

### 2.7. Subcutaneous Tumor Xenografts in Nude Mice

Mouse xenograft models were made by subcutaneous heterotransplantation of wild-type HepG2 (*WT*), *Nrf1α^−/−^*, *Nrf1α*-restored and *TCF11*-restored cells, respectively, into nude mice as described [51]. Each line of experimental cells (1 × 10^7^) were allowed for its exponential growth and then suspended in 0.2 mL of serum-free DMEM, before being inoculated subcutaneously into the right upper back region of male nude mice (BALB/C^nu/nu^, 6 weeks, 18 g, from HFK Bioscience, Beijing) at a single site. The procedure of injection into all mice was complete within 30 min, and subsequent formation of the subcutaneous tumor xenografts was observed. The tumor sizes, after emerged, were measured every two days, until the 42^nd^ day when all those mice were sacrificed and the transplanted tumors were excised. The sizes of growing tumors were calculated by a standard formula (i.e., V = ab^2^/2) and shown graphically (n = 7 per group). Notably, all the mice were maintained under standard animal housing conditions with a 12-h dark cycle and allowed access *ad libitum* to sterilized water and diet. All relevant studies were carried out on 8-week-old mice (with the license No. PIL60/13167) in accordance with United Kingdom Animal (Scientific Procedures) Act (1986) and the guidelines of the Animal Care and Use Committees of Chongqing University and the Third Military Medical University, both of which were also subjected to the local ethical review (in China). All relevant experimental protocols were approved by the University Laboratory Animal Welfare and Ethics Committee (with two institutional licenses SCXK-PLA-20120011 and SYXK-PLA-20120031). As for additional ethical concerns about the xenograft model mice bearing so big tumors insomuch as to give rise to certain bleeding ulcers, such a bad health condition of mice was only emerged from only day 2 prior to being sacrificed, and also such relevant study was indeed conducted according to the valid ethical regulations that have been approved.

### 2.8. The Colony Formation Assay on Soft Agar

The cell culture plates (each with a diameter of 10 cm) were coated by the basement gel containing 0.6% soft agar mixed in the complete medium, upon which the upper gel containing 0.35% soft agar. Then, experimental cells (2 × 10^4^, that had been growing in the exponential phase) was allowed for two-layer gel formation. Thereafter, the plates were cultured for 2-3 weeks in the incubator at 37°C with 5% CO_2_ before being stained with 1% crystal violet reagent (Sigma) and counted.

### 2.9. The in Vitro Scratch Assays

When experimental cells (1 × 10^5^) grown in 6-well plates reached 70% confluency, they were allowed for synchronization by 12-h starvation in serum-free medium and then treated with 1 μg/mL of mitomycin C (from Cayman, USA) for 6 h. Subsequently, a clear ‘scratch’ in the cell monolayer was created and then allowed for being healed in the continuous culture at 37°C with 5% CO_2_. Thereafter, the cell migration was quantified according to the standard procedures [52].

### 2.10. The Transwell-Based Migration and Invasion Assays

The transwell-based migration and invasion assays were conducted in the modified Boyden chambers (Transwell; Corning Inc. Lowell, MA, USA) as described previously [53]. When the growing cells reached 70% confluency, they were starved for 12 h in serum-free medium. The experimental cells (5 × 10^3^) were suspended in 0.5 mL medium containing 5% FBS and seeded in the upper chamber of a transwell, which allows the cells to grow on the microporous polycarbonate membrane that is tissue culture-treated to enhance the cellular attachment to the bottom. The cell-seeded transwells were placed in each well of 24-well plates containing 1 mL of complete medium (i.e., the lower chamber), and then cultured for 24 h in the incubator at 37°C with 5% CO_2_. Of note, the bottom of upper transwell was pre-coated by matrigel basement matrix (BD, Biosciences, USA), before the cells were placed in the invasion assay. The remaining cells in the upper chamber were removed, and the cells attached to the lower surface of the transwell membranes were fixed with 4% paraformaldehyde (AR10669, BOSTER) and stained with 1% crystal violet reagent (Sigma) before being counted.

### 2.11. Subcellular Fractionation

Equal numbers (4 × 10^6^) of different cell lines were seeded in each of 10-cm dishes and allowed for growth for 24 h, followed by treatment with Tet (1 μg/mL) for additional 12 h alone or in combination with MG132 (10 μmol/L, within the last four hours added) before being harvested in an ice-cold Nuclei EZ lysis buffer (Sigma, NUC101-1KT). Then the lysates were subjected to subcellular fractionation by centrifuging at 500 × *g* for 5 min at 4°C. The supernatants were collected as the non-nuclear cytoplasmic fractions, while the sediments were washed twice with the Nuclei EZ lysis buffer, and the resulting nuclear fractions were pelleted by centrifuging at 500 × *g* for 5 min at 4°C. Subsequently, the cytoplasmic and nuclear fractions were evaluated by Western blotting with distinct antibodies.

### 2.12. Immunofluorescence Assay

Experimental cells (2 × 10^5^) were allowed for 24-h growth on a cover glass placed in each of 6-well plates, and treated with Tet (1 μg/mL) for additional 12 h before being fixed with 4% paraformaldehyde for 20 min. The cells were then permeabilized for 10 min with 0.1% Triton X-100 (Beyotime, diluted with PBS) before immunocytochemistry with the primary antibodies against the V5 tag (diluted at 1:100) incubated at 4°C overnight. The immunostained cells were visualized by incubation with the fluorescein-conjugated goat anti-mouse IgG (ZSGB-BIO, dilution 1:100) for 1 h at room temperature in the dark, followed by DAPI staining (KeyGEN BioTECH, KGA215) of the nuclear DNAs for 5 min. The resulting images were observed and photographed by fluorescence microscope.

### 2.13. Flow Cytometry Analysis of Cell Cycle and Apoptosis

Experimental cells (5 × 10^5^) were allowed for growth in 6-cm dish for 48 h and synchronization by 12-h starvation in a serum-free medium, before being treated with 10 μmol/L BrdU for 12 h. The cells were fixed for 15 min with 100 μL BD Cytofix buffer (containing a mixture of the fixative paraformaldehyde and the detergent saponin) at room temperature and permeabilized for 10 min with 100 μL BD Cytoperm permeabilization buffer (containing fetal bovine serum as a staining enhancer) on ice. Subsequently, the cells were re-fixed and treated with 100 μL DNase (at a dose of 300 μg/mL in DPBS) for 1 h at 37°C, in order to expose the incorporated BrdU, followed by staining with FITC conjugated anti-BrdU antibody for 1 h at room temperature. Thereafter, the cells were suspended in 20 μL of 7-amino-actinomycin D solution 20 min for the DNA staining, and re-suspended in 0.5 mL of a staining buffer (i.e., 1 × DPBS containing 0.09% sodium azide and 3% heat-inactivated FBS), prior to the cell cycle analysis by flow cytometry. Additional fractions of cells (5 × 10^5^) were allowed for 48-h growth in 6-cm dish before being harvested for apoptosis analysis. The cells were pelleted by centrifuging at 500 × *g* for 5 min and washed by PBS three times, before being incubated for 15 min with 5 μL of Annexin V-FITC and 10 μL of propidium iodide (PI) in 195 μL of binding buffer, followed by apoptosis analysis with flow cytometry. The results were further analyzed by the FlowJo 7.6.1 sofware.

### 2.14. Hematoxylin-Eosin Staining Assay

Representatives of the above xenograft tumor tissues were fixed with 4% paraformaldehyde and then transferred to 70% ethanol according to the routine protocol. Thereafter, all individual tumor tissues were placed in the processing cassettes, dehydrated through a serial of alcohol gradient, and then embedded in paraffin wax blocks before being sectioned into a series of 5-μm-thick slides. Next, the tissue sections were dewaxed in xylene, and then washed twice in 100% ethanol to eliminate xylene, followed by rehydration in a series of gradient concentrations of ethanol with being distilled. Subsequently, they were stained with the standard hematoxylin and eosin (H&E) and visualized by microscopy.

### 2.15. Statistical Analysis

Statistical significances were determined using either Student’s *t*-test (for a comparison between two groups) or two-way ANOVA (for comparison among multiple groups). The relevant data presented herein are shown as a fold changes (mean ± SEM or ± SD) with significant differences that were calculated by the value of *p* < 0.05).

## 3. Results

### 3.1. Controllable Model Cell Lines for Stably Expressing TCF11, TCF11^ΔN^, Nrf1α, or Nrf2 are Established

As shown in Figure 1A, four distinct expression constructs for *TCF11*, *TCF11^ΔN^*, *Nrf1α* or *Nrf2*, together with a recombinase Flp-expressing plasmid, were co-transfected into the Flp-In™ T-REx™-293 host cells and also integrated into this host cells by Flp-mediated homologous recombination at FRT sites. Then, putative positive cell clones stably expressing the indicated gene of interest (GOI) were selected and maintained by hygromycin B (150 μg/mL) and blasticidin (15 μg/mL) to establish these cell models for expression of *TCF11*, *TCF11^ΔN^*, *Nrf1α* or *Nrf2*. Such model cell lines are controllable because their GOI are tightly monitored by interaction of Tet (tetracycline) with its repressor TetR; thus, only after TetR will be released from the TetO_2_ operator, transcriptional expression of GOI and down-stream target genes can be induced. In subsequent parallel experiments, an additional cell line co-transfected with the empty expression vector served as an internal negative control. In order to further validate such stable controllable expression of *TCF11*, *TCF11^ΔN^*, *Nrf1α* or *Nrf2,* respectively, all these indicated model cell lines were treated with 1 μg/mL Tet for 12 h and then determined by real-time quantitative PCR (Figure 1, B-D) and immunoblotting (Figure 1, E-G).

Notably, a pair of specific TCF11-recognized primers, including part of the Neh4L-coding nucleotides, were designed by distinguishing it from *Nrf1α*, because the Neh4L-missing nucleotides of *Nrf1α* remains present in *TCF11* and also in its N-terminally-truncated *TCF11^ΔN^*. Thus, the latter two factors-shared same primers were employed for quantitative PCR examinations of *TCF11* and *TCF11^ΔN^*. As anticipated, real-time qPCR revealed that each specific transcriptional expression of *TCF11*, *TCF11^ΔN^*, *Nrf1α* or *Nrf2,* respectively, was induced by Tet in their indicated model cell lines (Figure 1, B-D). Such Tet-induced protein expression abundances of TCF11, TCF11^ΔN^, Nrf1α and Nrf2 were evaluated by immunoblotting with antibodies against Nrf1/TCF11, Nrf2 and their C-terminally-tagged V5 epitope, respectively. The resulting data (Figure 1, E-G) demonstrated that those model cell lines had a strong capability to stably express each of interested genes, one of which had almost no effects on all the others examined, though each factor-specific expression was induced under Tet control. Of note, a major protein of TCF11 exhibited a slightly slower mobility, than that of Nrf1α, on electrophoretic gels, whereas the V5-tagged TCF11^ΔN^ mobility appeared to coincide closely to the electrophoretic band of Nrf2 (Figure 1E). Besides, a few of putative C-terminally-truncated isoforms of Nrf1α, TCF11 or TCF11^ΔN^ were also immunoblotted with Nrf1/TCF11-specfic antibody (Figure 1F). Furtherly, the subcellular nucleocytoplasmic fractionation and immunofluorescence experiments unraveled that a certain amount of TCF11, TCF11^ΔN^, Nrf1α or Nrf2 was allowed to be localized in the cellular nucleus (Figure S3, A-E). However, such nuclearly-positioning proteins are rapidly degraded, due to this fact that these protein degradations were inhibited by MG132 (at 10 μmol/L), so that obvious increases in their protein expression levels were recovered in the nuclear fractions of MG132-treated cells (Figure S3, A-D).

For further time-course analysis of *TCF11*, *TCF11^ΔN^*, *Nrf1α* and *Nrf2*, their indicated cell lines that had been pre-treated for 12 h with 1 μg/mL Tet were treated with 50 μg/mL cycloheximide (CHX, that inhibits biosynthesis of nascent proteins) alone or plus a proteasomal inhibitor MG132. As shown in Figure 1 (H-K), the N-terminal truncation of TCF11 (to remove both its ER-targeting signal peptide sequence and adjacent juxtamembrane proteolytic degron) caused the resulting TCF11^ΔN^ isoform to become stabilized relatively. Subsequent calculation of their half-lives in CHX-treated cells suggested that a major TCF11^ΔN^ protein was conferred with a relative higher stability than those of TCF11, Nrf1 and Nrf2, each with distinct expression isoforms (as illustrated graphically in Figure 1, H-K). Upon co-treatment of cells with CHX and MG132, all those examined protein half-lives were markedly enhanced. Such being the case, intact TCF11 protein-A became gradually fainter and then disappeared by 2 h after co-treatment (Figure 1H). Such disappearance of TCF11 protein-A seemed to be accompanied by gradual emergence and increment of its protein-B and -C until the end of 8-h experimentation. Similar yet different conversion of Nrf1α protein-A into short isoforms-B, -C and -D was observed (Figure 1I). However, no similar changes were determined in two cases of Nrf2 and TCF11^ΔN^, because both proteins gradually decreased with increasing time of co-treatment until they finally disappeared from 4 h to 8 h after co-treatment of the cells with CHX and MG132 (Figure 1, J & K). Collectively, these demonstrate that the absence of the N-terminally ER-targeting sequence in TCF11^ΔN^, as well as in Nrf2, allows them to display distinguishable behaviors from the ER-resident TCF11 and Nrf1α, both of which are manifested in similar but nuanced ways of topobiological processing within and around this organelle before being dislocated into the nucleus.

### 3.2. Differential Expression Profiles of Genes Regulated by TCF11, TCF11^ΔN^, Nrf1α and Nrf2 are Defined

To identify differential expression profiles of genes regulated by TCF11, TCF11^ΔN^, Nrf1α and/or Nrf2, relevant RNAs extracted from the established model cell lines were subjected to transcriptome sequencing. As a result, all those detectable genes, if upregulated or downregulated respectively with fold changes ≥ 2 or ≤ 0.5 plus false discovery rate (FDR) ≤ 0.001 (Figure 2A), were defined as differentially expressed genes (DEGs), by comparison with equivalents measured from control cells. Thereof, 2845 DEGs were detected in *TCF11-*expressing cells, of which 2786 target genes were upregulated by TCF11 (Figure 2A, and Table S2). By contrast, *Nrf1α-*expressing cells only yielded 1001 DEGs, i.e., 957 upregulated plus 44 downregulated (Table S3), whereas *Nrf2-*expressing cells led to a significantly decreased number of upregulated genes (i.e., 276) but as accompanied by down-regulation of 457 DEGs (Table S4). Notably, 1459 DEGs were identified in *TCF11^ΔN^-*expressing cells, with so many as 989 genes downregulated (Table S5). Interestingly, TCF11^ΔN^ appeared to be endowed with a regulatory trend of its target genes similar to that of Nrf2 (Figure 2). Such changed DEGs with distinct trends among different groups were further explicated by scatterplots of gene expression profiles (Figure S4A). Together, these data indicate that TCF11 makes a greater impact on the overall gene expression than Nrf1α, albeit both contribute to basically positive regulation of their DEGs, whereas Nrf2 and TCF11^ΔN^ make more contributions to negative regulation rather than positive regulation of their DEGs.

**Figure 2.**
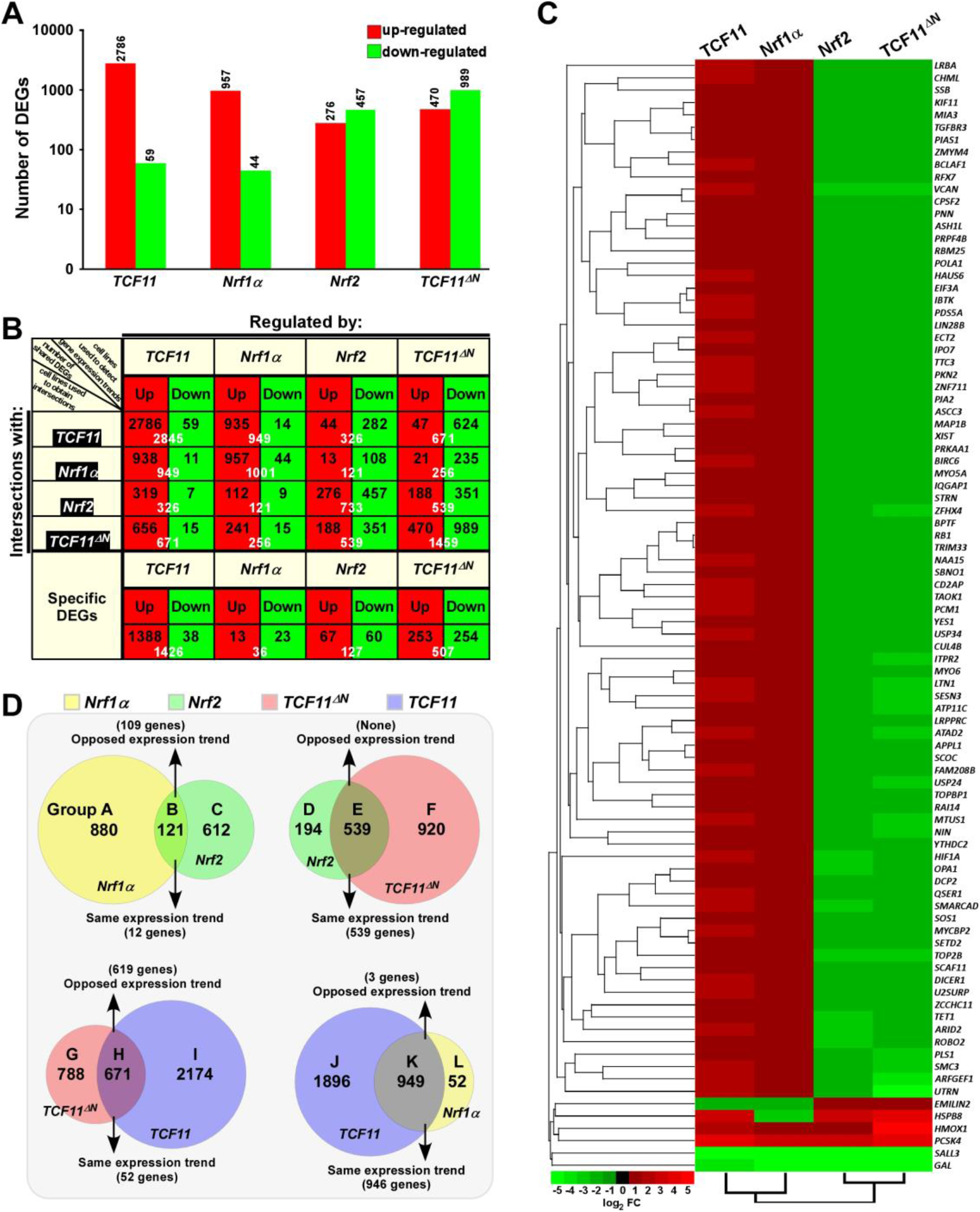
Statistical analysis of the data obtained from transcriptome sequencing. (**A**) Differentially expressed genes (DEGs) in distinct cell lines were analyzed by transcriptome sequencing, and relevant differences in the number of those increased or decreased DEGs are shown in the histogram. The DEGs were selected according to the following criteria: fold change ≥ 2 or ≤ 0.5 and FDR ≤ 0.001 (as compared to the Control cells). (**B**) The specific DEGs in each cell line and their common DEGs between every two cell lines were also counted as indicated in the chart, and the number of increased and decreased DEGs in each group is shown separately in black font, and the total is shown in white. In addition, the change trends of DEGs in each group were indicated in red or green, which represent up-regulated or down-regulated in the cells in the first row, respectively. (**C**) The heatmap with hierarchical clustering of 90 DEGs shared in all four cells lines. (**D**) Distinct groupings of the subsequent functional annotation and also the Venn diagram of DEGs between every two cell lines.

The intersections of these four groups of DEGs regulated by TCF11, TCF11^ΔN^, Nrf1α and/or Nrf2 were shown by the Venn diagram (Figure S4B). Those common DEGs between every two cell groups and each specific DEGs were taken into account (in *orthogonal table*, Figure 2B), with distinct regulatory tendencies of DEGs even in each subgroup. Of note, the common DEGs between *Nrf2-*expressing cells and the others were more likely to be downregulated rather than up-regulated by Nrf2, with a roughly similar number of its upregulated genes to down-regulated genes amongst *Nrf2*-specific DEGs. Another similar situation also occurred in *TCF11^ΔN^-*expressing cells.

Approximately 90 DEGs were identified to be shared among these four cell lines (as shown in the Venn diagram, Figure S4B). Differential expression levels of these DEGs were presented by their heatmap with hierarchical clustering (Figure 2C). In the shared DEGs, only four genes *HMOX1* (heme oxygenase 1), *PCSK4* (proprotein convertase subtilisin/kexin type 4), *SALL3* (spalt like transcription factor 3) and *GAL* (galanin and GMAP prepropeptide) had a similar expression trends in all four cell lines. Most of other shared DEGs were up-regulated by TCF11 and Nrf1α, but down-regulated by Nrf2 and TCF11^ΔN^, besides two exceptions of *EMILIN2* (elastin microfibril interface 2) and *HSPB8* (heat shock protein family B member 8) regulated by opposite ways (Figure 2C). In addition, no evident effects of *Nrf2-*expressing cell model on *TCF11*, *TCF11^ΔN^* or *Nrf1α* was examined by transcriptome sequencing, but conversely, only a marginal increase of *Nrf2* expression was found in *Nrf1α-*, *TCF11-*, rather than *TCF11^ΔN^-*expressing cell models (Figure S4C). Amongst other CNC-bZIP members, only *BACH1* was up-regulated by TCF11, whilst all *sMAF* partners were up-regulated by Nrf2 and TCF11^ΔN^, but *Keap1* was unaffected.

In order to gain a further insight into similarities and differences in biological functions of between TCF11, TCF11^ΔN^, Nrf1α and Nrf2, their common and different regulatory DEGs were scrutinized in distinct combinations of every two groups as indicated by Groups A to L (Figure 2D). Similar or opposite trends in those common DEGs of every two cell groups were schematically shown. For an example of Group B, 121 common DEGs between *Nrf1α* and *Nrf2* were subdivided into 109 oppositely-regulated genes and the other 12 genes with the same directional tend. In Group E, all those common DEGs were manifested only with the same directional trend to be regulated by both *Nrf2* and *TCF11^ΔN^*. By contrast, a largely opposing expression trend in 619 of the common 671 DEGs co-regulated by *TCF11^ΔN^* and *TCF11* was shown in Group H, whilst 946 of the other common 949 DEGs shared by *TCF11* and *Nrf1α* in Groups K showed the same tendency of expression change. Such groups of these DEGs were also subjected to comprehensive analysis of their functional annotations as described below.

### 3.3. Nrf1α and Nrf2 Have Diverse Regulatory Patterns of Their Target Genes

In Group A, 880 DEGs were identified in *Nrf1α*-expressing rather than *Nrf2*-expressing cells, and further subjected to their functional annotation by the DAVID (*<u>d</u>*atabase for *<u>a</u>*nnotation, *<u>v</u>*isualization and *<u>i</u>*ntegrated *<u>d</u>*iscovery) (Figure 3A), for the data mining in order to delineate unique biological functionality of Nrf1α in regulating genes preferentially than Nrf2. Besides their shared common 121 DEGs in Group B, the other 612 DEGs in Group C were also annotated to identify those biological functions of Nrf2 that were responsible preferentially than Nrf1α (Figure 3A, and Table S6). The details of all top significant biological process terms and pathways enriched by DEGs in three different groups were deciphered in histograms and scatterplots (Figure 3A). Furtherly, the biological process terms and pathways were classified by using QuickGO and KEGG databases. Such functional annotation analysis implied that DEGs regulated by Nrf1α (Group A) were predominantly involved in cellular metabolic process, response to stimulus, replication and repair, cell growth and death, protein folding, sorting and degradation, signal transduction, immune system and cancers. In Group B, DEGs co-regulated by Nrf1α and Nrf2 took part in distinct cellular metabolic process, RNA processing, regulation of biological process, response to stimulus, apoptotic process, regulation of transcription, translation and cancers. Nrf2-regulated DEGs (in Group C) were also responsible for cellular metabolic process, apoptotic process, response to stimulus, regulation of transcription, development and regeneration, signal transduction, endocrine system, infectious diseases and cancers.

**Figure 3.**
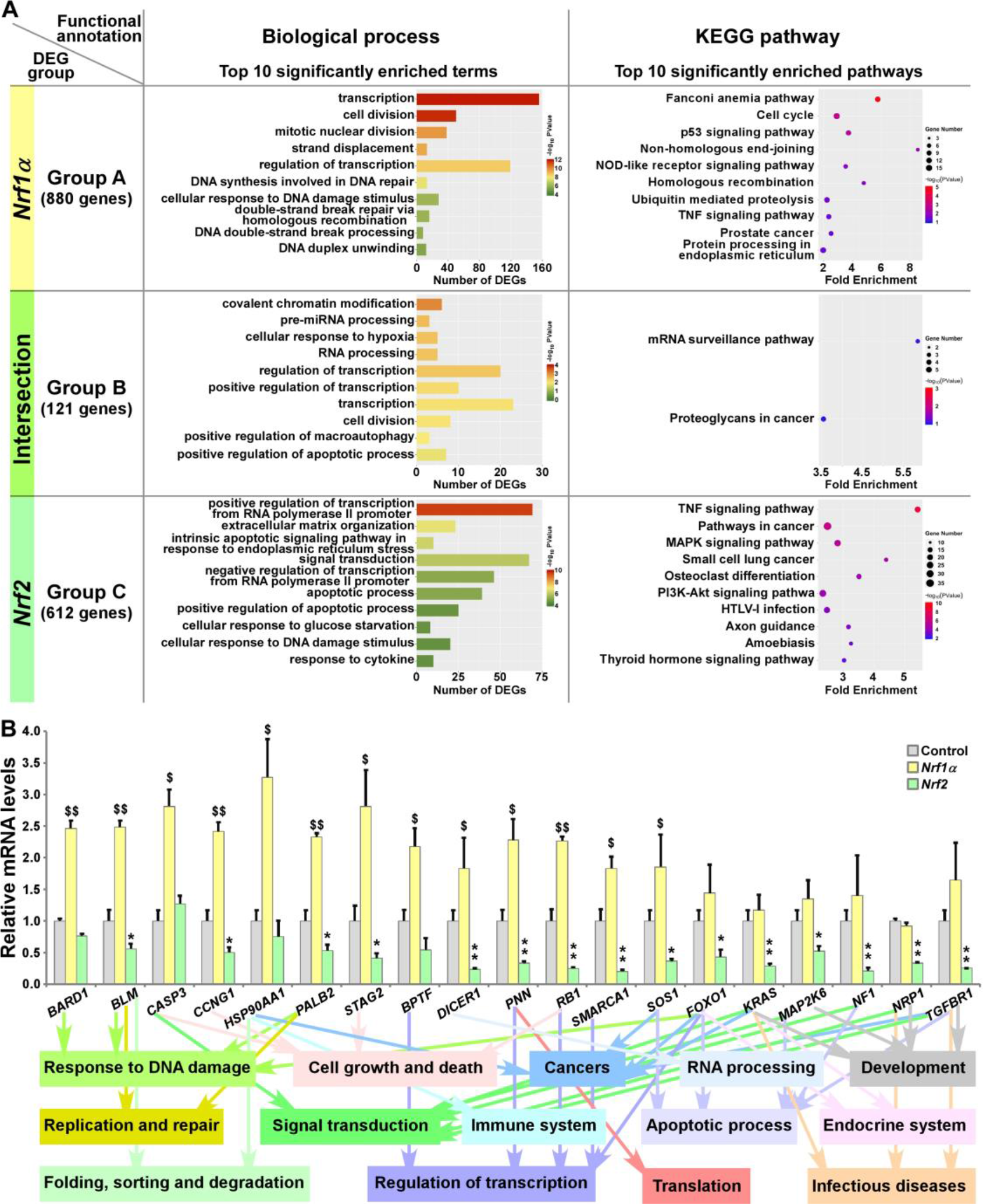
Functional annotation of specific or common DEGs in *Nrf1α* and *Nrf2* cells. (**A**) The top 10 of significant biological process terms and pathways enriched by DEGs in Groups A, B, and C were exhibited in histograms and scatterplots, respectively. (**B**) After *Nrf1α*, *Nrf2* and Control cell lines were incubated with 1 μg/mL Tet for 12 h, total RNAs were isolated and reversely transcribed into the first strand of cDNA. Subsequently, the mRNA levels of DEGs that were associated with more functions, along with high expression levels and well significance in Groups A to C, were determined by quantitative real-time PCR analysis of *Nrf1α*, *Nrf2* and Control cell lines. The data are shown as mean ± SEM (n = 3 × 3, * *p* < 0.05; ** *p* < 0.01; $, *p* < 0.05; $$, *p* < 0.01, when compared to the *Control values*).

Based on certain association with multiple functions, along with higher expression levels and well significance, 19 DEGs were selectively verified by further quantitative PCR analysis (Figure 3B). The results demonstrated that 4 DEGs, including *BARD1* (*BRCA1 associated RING domain 1*), *CASP3* (*caspase 3*), *HSP90AA1* (*heat shock protein 90 alpha family class A member 1*) and *BPTF* (*bromodomain PHD finger transcription factor*) were upregulated by Nrf1α, but not significantly affected by Nrf2. By contrast, 9 DEGs, including *BLM* (*Bloom syndrome*, *RecQ like helicase*), *CCNG1* (*cyclin G1*), *PALB2* (*partner and localizer of BRCA2*), *STAG2* (*stromal antigen 2*), *DICER1* (*dicer 1*, *ribonuclease III*), *PNN* (*pinin*, d*esmosome associated protein*), *RB1* (*RB transcriptional corepressor 1*), *SMARCA1* (*SWI/SNF related*, *matrix associated*, *actin dependent regulator of chromatin*, *subfamily a*, *member 1*) and *SOS1* (*SOS Ras/Rac guanine nucleotide exchange factor 1*) were upregulated by Nrf1α, but downregulated by Nrf2. Another 6 DEGs, such as *FOXO1* (f*orkhead box O1*), *KRAS* (*KRAS proto-oncogene*, *GTPase*), *MAP2K6* (*mitogen-activated protein kinase kinase 6*), *NF1* (*neurofibromin 1*), *NRP1* (*neuropilin 1*) and *TGFBR1* (*transforming growth factor beta receptor 1*) were reduced in *Nrf2-*expressing cells but, but no significant changes of them was detected in *Nrf1α-*expressing cells. In addition, putative functions of the examined genes as well as of their encoding proteins were also extracted (Figure 3B, *on the bottom*).

### 3.4. TCF11^ΔN^ Exhibits a Similar Regulatory Profile to that of Nrf2

All those DEGs regulated by *Nrf2* or *TCF11^ΔN^* alone or both were divided into three Groups D, E and F, respectively, and then functionally annotated with the above-described methods (Figure 4A and Table S7). In Group D, 194 DEGs regulated by Nrf2, but not by TCF11^ΔN^, were generally involved in cellular metabolic process, localization, regulation of biological process, developmental process, response to stimulus, signal transduction and cancers. In the intersected Group E, 539 DEGs co-regulated by *Nrf2* and *TCF11^ΔN^* were also associated with cellular metabolic process, development and regeneration, regulation of transcription, response to stimulus, apoptotic process, endocrine system, signal transduction, infectious diseases and cancers. In Group F, 920 DEGs regulated by *TCF11^ΔN^* were significantly enriched in cellular metabolic process, cell motility, cellular community, developmental process, regulation of transcription, signal transduction, signaling molecules and interaction, cardiovascular diseases and cancers. Intriguingly, 8 common biological process terms and additional 8 common pathways were predicted to exist between the top 10 functions significantly enriched in Group C (including DEGs regulated by *Nrf2* but not by Nrf1α) and Group E (with an intersection of DEGs shared by *Nrf2* and *TCF11^ΔN^*) (Figure S4D). This implies that Nrf2 and TCF11^ΔN^ share the common regulatory profiles, but they are likely differential from those of Nrf1α and Nrf2.

**Figure 4.**
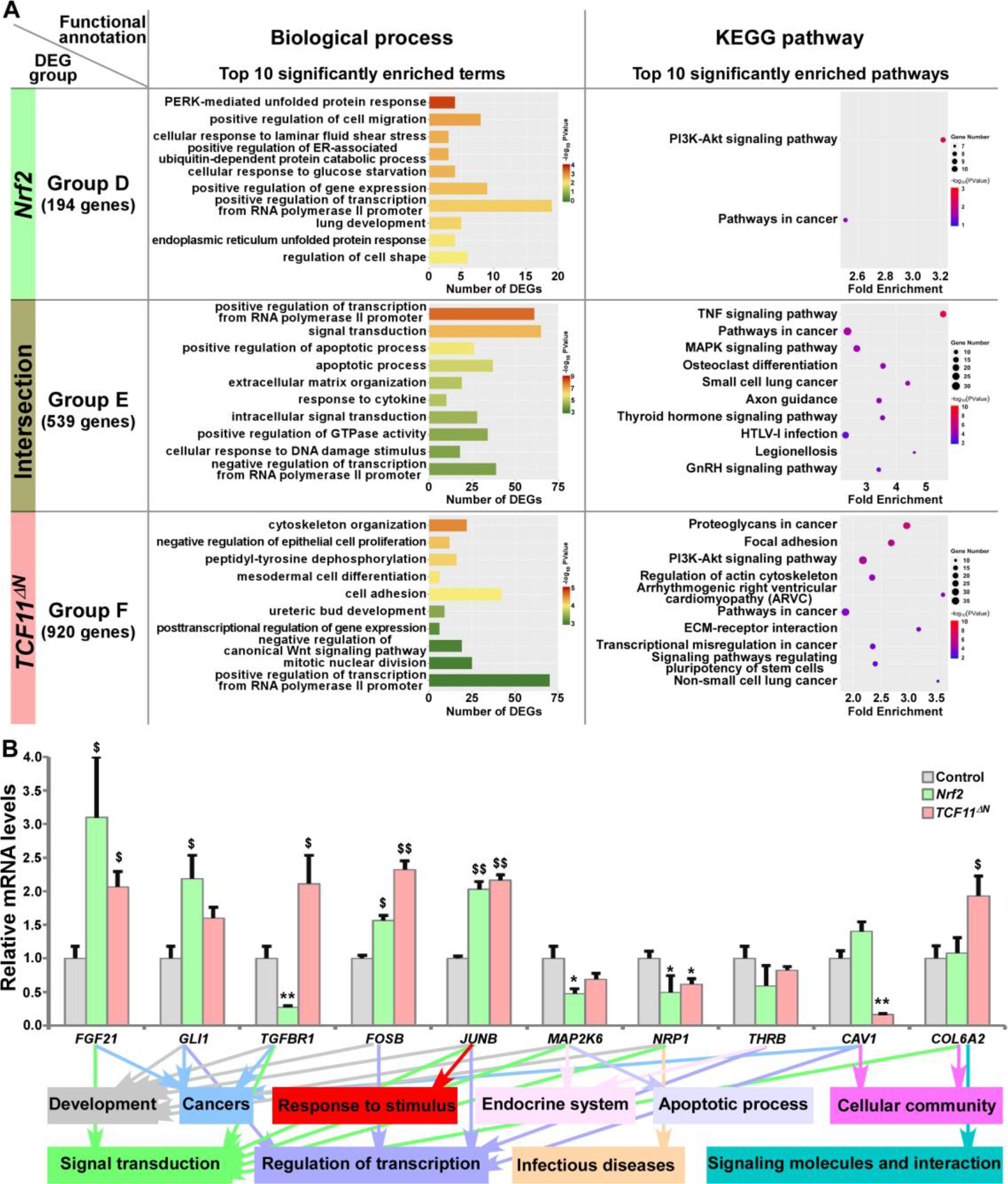
Functional annotation of specific or common DEGs in *Nrf2* and *TCF11^ΔN^* cells. (**A**) Top 10 of significant biological process terms and pathways enriched by DEGs in Groups D, E, and F were exhibited in histograms and scatterplots, respectively. (**B**) After induced with 1 μg/mL Tet for 12 h, total RNAs were isolated from *Nrf2*, *TCF11^ΔN^* or Control cell lines and then reversely transcribed into the first strand of cDNA. Subsequently, the mRNA levels of DEGs that were associated with more functions as annotated, along with high expression levels and well significance in Groups D to F, were determined by quantitative real-time PCR analysis of *Nrf2*, *TCF11^ΔN^* and Control cells. The data are shown as mean ± SEM (n = 3 × 3, * *p* < 0.05; ** *p* < 0.01; $, *p* < 0.05; $$, *p* < 0.01, when compared to the *Control values*).

The results of quantitative PCR validation (Figure 4B) revealed that expression of *FGF21* (*fibroblast growth factor 21*), *FOSB* (a *subunit of AP-1 transcription factor*) and *JUNB* (*another subunit* of *AP-1*) were upregulated by Nrf2 and TCF11^ΔN^, with downregulation of *MAP2K6* (*mitogen-activated protein kinase kinase 6*) and *NRP1* (*neuropilin 1*). By contrast, *GLI1* (*GLI family zinc finger 1*) was upregulated by Nrf2, but not significantly altered by TCF11^ΔN^. Conversely, *TGFBR1* (*transforming growth factor beta receptor 1*) was downregulated by Nrf2, but upregulated by TCF11^ΔN^. However, reduced expression of *CAV1* (*caveolin 1*) was accompanied by increased *COL6A2* (*collagen type VI alpha 2 chain*) in TCF11^ΔN^-expressing cells, but almost unaffected by Nrf2. In addition, putative functions of such target genes were mapped, as indicated by the histogram (Figure 4B, *on the bottom*).

### 3.5. TCF11 and Its Truncated TCF11^ΔN^ Regulate Similar yet Different Subsets of Target Genes

The common and distinct target DEGs in *TCF11^ΔN^*- and/or *TCF11*-expressing cell lines were assigned into three groups, and functionally annotated with the aforementioned method as visualized in bar charts and scatterplots (Figure 5A, and Table S8). In Group G, 788 DEGs were identified as targets of TCF11^ΔN^, but not of TCF11, and enriched with distinct functions in cellular metabolic process, regulation of biological process, cellular community, apoptotic process, developmental process, development and regeneration, response to stimulus, signal transduction, signaling molecules and interaction, infectious diseases and cancers. Their commonly-shared 671 DEGs in Group H were functionally responsible for cellular metabolic process, cell cycle, cell motility, signal transduction; protein folding, sorting and degradation, transport and catabolism, endocrine system and cancers. In Group I, 2174 DEGs were regulated by *TCF11*, rather than by *TCF11^ΔN^*, and preferentially functionally associated with cellular metabolic process, cell growth and death, replication and repair, response to stimulus, signal transduction; protein folding, sorting and degradation, regulation of transcription, translation, transport and catabolism, carbohydrate metabolism and infectious diseases. Notably, 6 identical pathways were found by comparison between top 10 significantly enriched pathways from Group C (i.e., DEGs regulated by *Nrf2* but not by Nrf1α) and Group G (i.e., DEGs regulated by *TCF11^ΔN^* but not by TCF11) (Figure S4D). This indicates that TCF11^ΔN^-regulated genes are much likely to execute somewhat combinational or overlapping functions with Nrf2-target genes.

**Figure 5.**
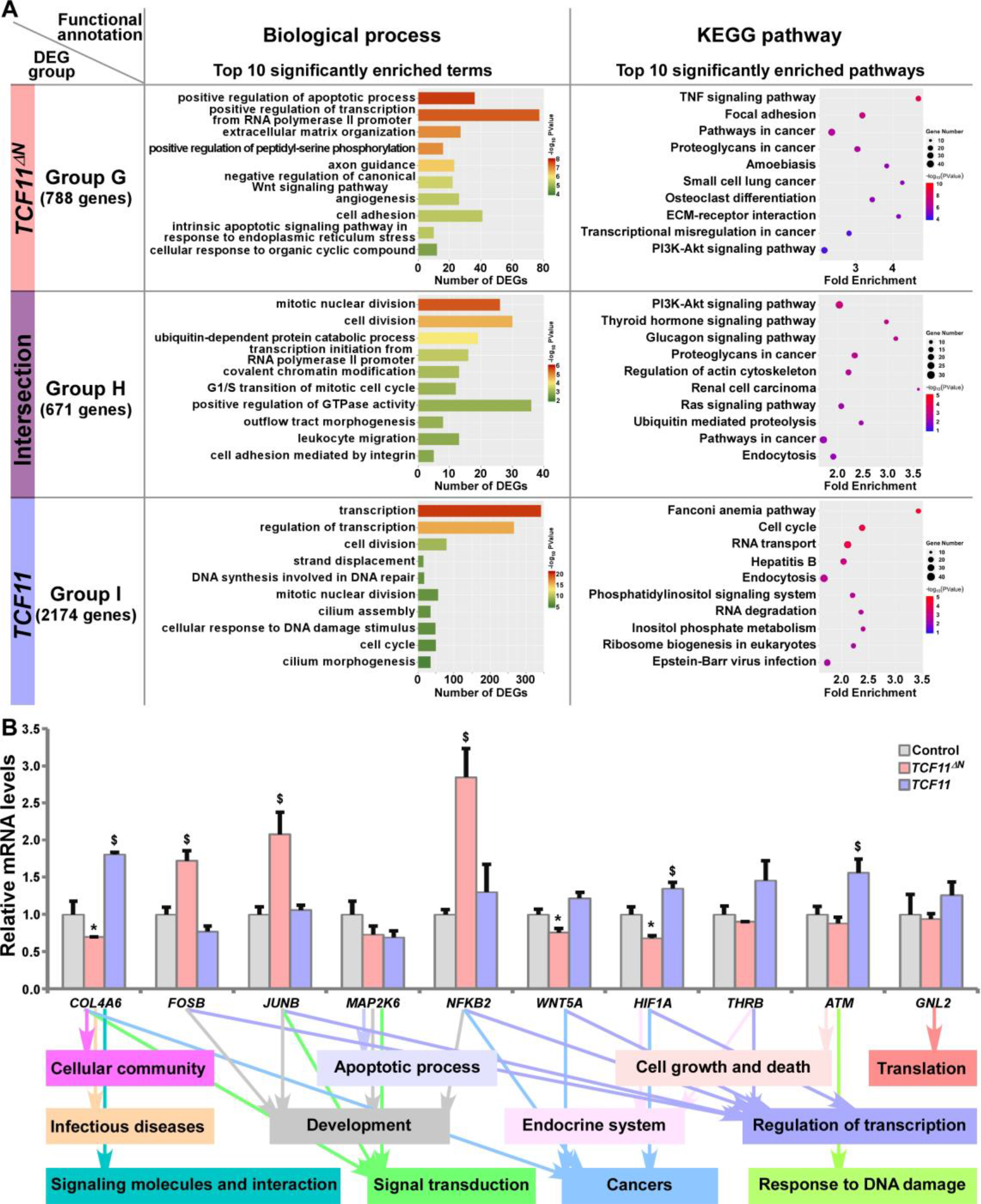
Functional annotation of specific or common DEGs in *TCF11^ΔN^* and *TCF11* cells. (**A**) Top 10 of significant biological process terms and pathways enriched by DEGs in Groups G, H, and I were exhibited in histograms and scatterplots, respectively. (**B**) After induced with 1 μg/mL Tet for 12 h, total RNAs were isolated from Control, *TCF11^ΔN^* or *TCF11* cell lines before being reversely transcribed into the first strand of cDNA. Subsequently, relative mRNA levels of DEGs that were associated with more functions, along with high expression levels and well significance in Groups G to I, were determined by quantitative real-time PCR in Control, *TCF11^ΔN^* and *TCF11* cells. The data are shown as mean ± SEM (n = 3 × 3, * *p* < 0.05; $, *p* < 0.05, when compared to the *Control values*).

Subsequently, several unique or common target genes regulated by TCF11 and/or TCF11^ΔN^ were also validated by quantitative PCR (Figure 5B). The results showed that *FOSB*, *JUNB* and *NFKB2* (*nuclear factor kappa B, subunit 2*) were upregulated, while *WNT5A* (*Wnt family member 5A*) was downregulated, by TCF11^ΔN^, but unaffected by TCF11. Conversely, *COL4A6* (*collagen type IV alpha 6*) and *HIF1A* (*hypoxia inducible factor 1 alpha*) were upregulated by TCF11, but downregulated by TCF11^ΔN^. Besides, *ATM* (*ATM serine/threonine kinase*) was also upregulated by TCF11, but roughly unaltered by TCF11^ΔN^. The putative functions relative to these examined genes were exhibited (as shown in Figure 5B, *on the bottom*).

### 3.6. TCF11 and Nrf1α Display Similar but Differential Regulatory Profiles

Those DEGs regulated by TCF11 and/or Nrf1α were grouped by J to L, and then functionally annotated by DAVID, with histograms and scatterplots exhibited (Figure 6A, and Table S9). Comprehensive analysis of the top significantly enriched biological process terms and pathways showed that 1896 DEGs of Group J, by identifying *TCF11-*, but not *Nrf1α*-, expressing cells, were associated with distinct functions in cellular metabolic process, cell cycle, subcellular localization, transport and catabolism, carbohydrate metabolism, regulation of transcription, translation, signal transduction, endocrine system, development and regeneration, infectious diseases and cancers. In Group K, TCF11 and Nrf1α co-regulated 949 DEGs that were involved in cellular metabolic process, cell growth and death, replication and repair, folding, sorting and degradation, regulation of transcription, response to stimulus, endocrine system, immune system and cancers. In Group L, only 52 DEGs were identified by transcriptional regulation by Nrf1α, but unaffected by TCF11 expression. Their putative functions were associated with cellular community and metabolic process, and subcellular localization, regulation of biological process, signal transduction and response to stimulus. Besides, many of overlapping functions were predicted to exist among distinct combinations of top significantly enriched functions exerted by Group A (i.e., DEGs regulated by Nrf1α but not by Nrf2), Group I (i.e., DEGs regulated TCF11 but not by TCF11^ΔN^) and Group K (i.e., DEGs co-regulated by TCF11 and Nrf1α) (Figure S4D). Thus, it is inferable that TCF11 and Nrf1α displays similar regulatory profiles, with a striking disparity from those of TCF11^ΔN^ or Nrf2.

**Figure 6.**
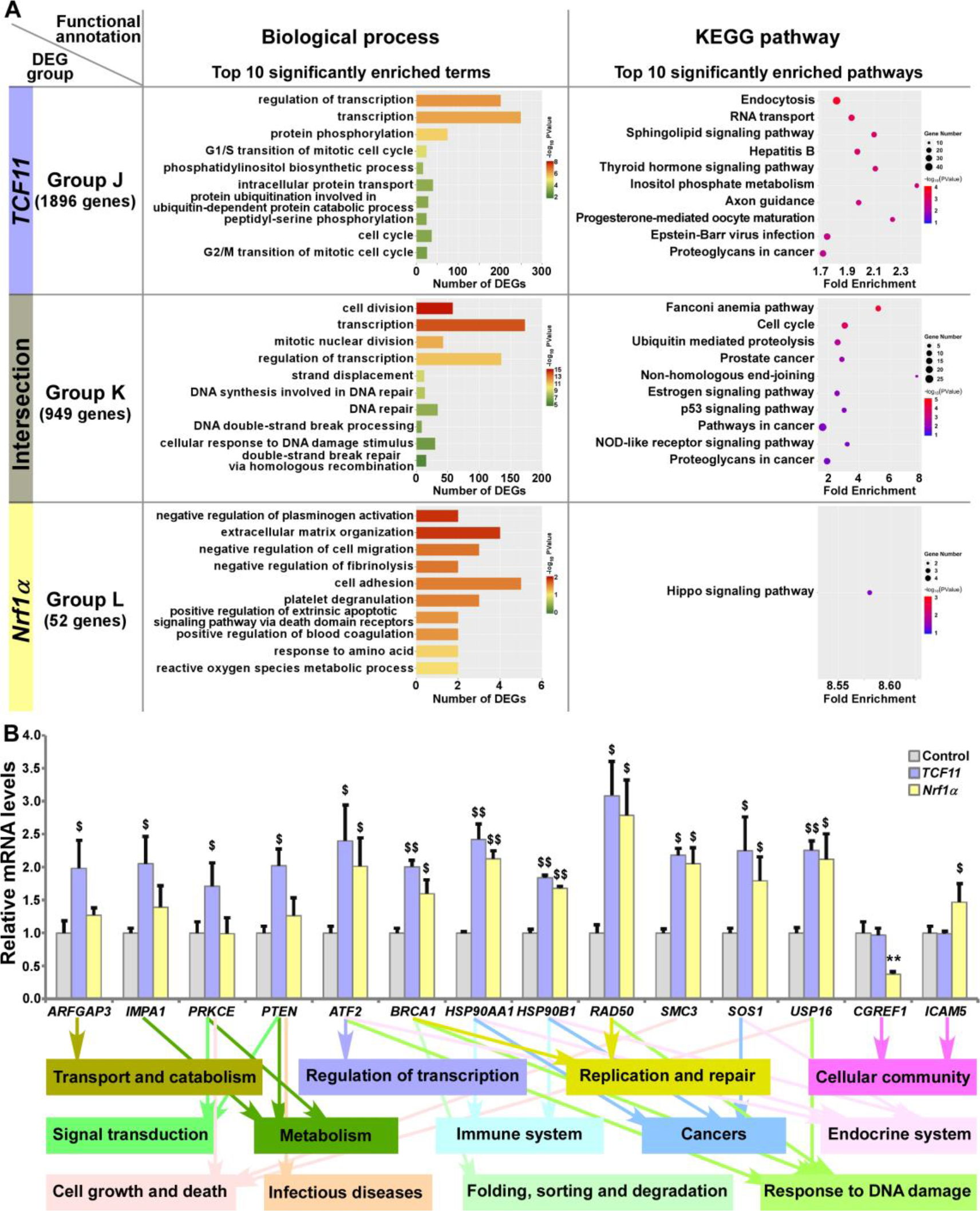
Functional annotation of specific or common DEGs in *TCF11* and *Nrf1α* cells. (**A**) Top 10 of significant biological process terms and pathways enriched by DEGs in Groups J, K, and L were exhibited in histograms and scatterplots, respectively. (**B**) After induced with 1 μg/mL Tet for 12 h, total RNAs were isolated from Control, *TCF11* or *Nrf1α* cell lines and then reversely transcribed into the first strand of cDNA. Subsequently, relevant mRNA levels of DEGs that were associated with more functions as annotated, along with high expression levels and well significance in Groups J to L, were determined by quantitative real-time PCR in Control, *TCF11* and *Nrf1α* cells. The data are shown as mean ± SEM (n = 3 × 3, ** *p* < 0.01; $, *p* < 0.05; $$, *p* < 0.01, when compared to the *Control values*).

Amongst the above DEGs, 14 were selected for further quantitation by real-time PCR (Figure 6B). The results revealed that 8 DEGs were upregulated by both TCF11 and Nrf1α, which included *ATF2* (*activating transcription factor 2*), *BRCA1* (*BRCA1 DNA repair associated*), *HSP90AA1* (*heat shock protein 90 alpha family class A member 1*), *HSP90B1* (*heat shock protein 90 beta family member 1*), *RAD50* (*RAD50 double strand break repair protein*), *SMC3* (*structural maintenance of chromosomes 3*), *SOS1* (*SOS Ras/Rac guanine nucleotide exchange factor 1*) and *USP16* (*ubiquitin specific peptidase 16*). Additional 4 DEGs, i.e., *ARFGAP3* (*ADP ribosylation factor GTPase activating protein 3*), *IMPA1* (*inositol monophosphatase 1*), *PRKCE* (*protein kinase C epsilon*) and *PTEN* (*phosphatase* and *tensin homolog*) were upregulated by TCF11, but almost unaffected by Nrf1α. Conversely, *Nrf1α* expression caused a decrease in *CGREF1* (*cell growth regulator with EF-hand domain 1*) as accompanied by increased *ICAM5* (*intercellular adhesion molecule 5*), but both genes were unaltered by TCF11. These examined genes were responsible for their putative functions as indicated (Figure 6B, *on the bottom*). Altogether, TCF11 and Nrf1α exhibit a similar regulatory profile, but with some of quietly different target genes.

Notably, the relative expression levels of those representative genes from Groups A to L in real-time quantitative PCR are basically consistent with the sequencing data, all with significant positive correlations (as shown in Figure S5). Collectively, the overall mapping profile between each of these four transcription factors and the enriched biological functions had been constructed according to the functional annotation and comparative analysis of each group of DEGs (Figure S6).

### 3.7. TCF11 is a more Potent Player than Nrf1α at Preventing Tumor Xenografts in Nude Mice

As analyzed above, distinct subset of DEGs regulated by Nrf1, TCF11 and/or Nrf2 were annotated for their functional relevancies to cancer development or prevention. In fact, our previous work had revealed that knockdown of Nrf1 caused a significant malignant growth of subcutaneous tumor xenografts in nude mice [54]. Thereof, Nrf1α was indicated to act as a dominant tumor-repressor insomuch as to confine oncogenicity of Nrf2 [16,17]. Herein, to corroborate the putative tumor-preventing effects of Nrf1α and TCF11, both CNC-bZIP factors were restored by transfecting the lentivirus expression constructs into HepG2 cells with a specific loss of *Nrf1α*, respectively, as described elsewhere [16]. As a consequence, the resulting *Nrf1α-* or *TCF11*-restored cell lines were confirmed by quantitative PCR and immunoblotting to be definitely true (Figure 7A).

**Figure 7.**
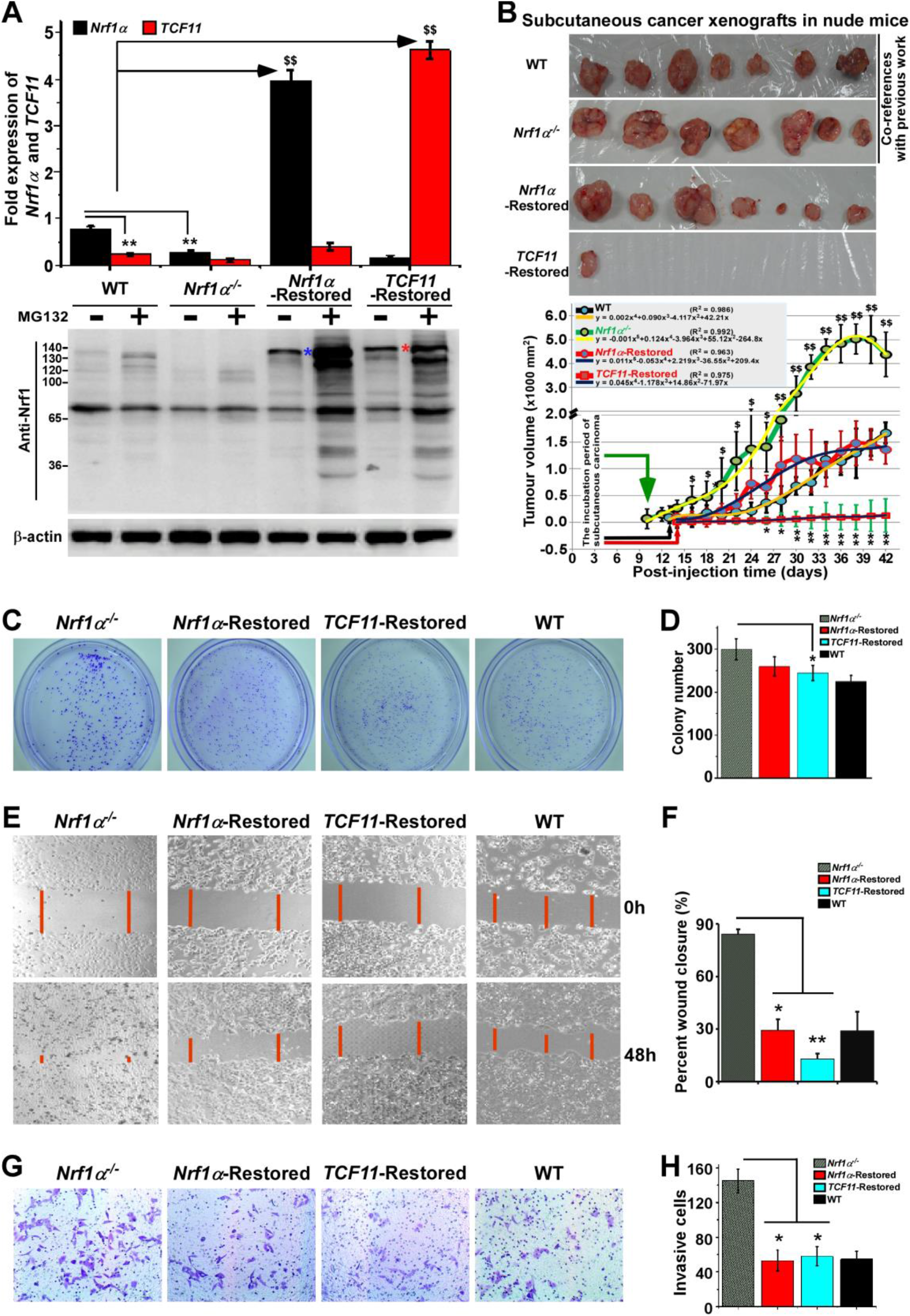
Malgrowth of *Nrf1α^−/−^*-derived hepatoma cells were significantly suppressed by restoration of *Nrf1α* and *TCF11*. (**A**) Both quantitative real-time PCR (*up*) and Western blotting (*down*) were employed to identify the protein and mRNA levels of Nrf1α and TCF11 in *Nrf1α*- and *TCF11*-Restored hepatoma cells. The experimental cells had been treated with or without 10 μmol/L MG132 for 4 h before being harvested for Western blotting. The data are shown as mean ± SEM (n = 3 × 3, ** *p* < 0.01; $$, *p* < 0.01). (**B**) Differences in mouse subcutaneous xenograft tumors derived from wild type HepG2 (WT), *Nrf1α*^−/−^, *Nrf1α*-Restored and *TCF11*-Restored cells were measured in size every two days, before being sacrificed on the 42^nd^ day. The data are shown as mean ± SD (n = 7 per group, * *p* < 0.05; ** *p* < 0.01; $, *p* < 0.05; $$, *p* < 0.01, when compared with the *WT* group). (**C-H**) The soft agar colony formation (C, D), as well as migration (E, F) and invasion (G, H), of wild type HepG2 (WT), *Nrf1α*^−/−^, *Nrf1α*-Restored and *TCF11*-Restored cells were examined as described in ‘Materials and Methods’. The data are shown as mean ± SD (n = 9, * *p* < 0.05, ** *p* < 0.01).

Subsequently, both *Nrf1α-* and *TCF11*-restored cell lines, alongside with *Nrf1α^−/−^* cells and wild-type (*WT*) HepG2 cells, were heterotransplanted into distinct groups of those immunodeficient nude mice at their subcutaneous loci as indicated. After tumor formation, the sizes of growing tumors were measured for every two days within ensuing five weeks before the tumor-bearing mice were sacrificed. The resulting data were calculated and shown graphically (Figure 7B), as a consequence demonstrating that restoration of *Nrf1α* or *TCF11* enables a significant tumor-preventing effect on the subcutaneous human carcinoma xenografts in nude mice, when compared with those obtained from *Nrf1α^−/−^* and WT cell lines. In addition, it should be also noted that both *Nrf1α^−/−^* and WT hepatoma cell lines, as reported previously [16], were herein used as only two co-references in this parallel animal experiments to strengthen their comparability (Figure 7B). Furthermore, a series of comparative experiments revealed that *Nrf1α^−/−^* cell proliferation, migration and invasion were all significantly suppressed by restoration of *Nrf1α* and *TCF11* (Figure 7, C-H), and the cell-cycle arrest at S-phase, along with the increase in early apoptosis, due to the restoration of them (Figure S7). This is further supported by pathohistological results of the hematoxylin and eosin (HE) staining, revealing that malignant progression of *Nrf1α^−/−^* -derived tumor xenografts was substantially suppressed by restoration of either *Nrf1α* or *TCF11* with complete coagulative necrosis of tumor tissues (Figure S8). Collectively, TCF11 acts as a more potent tumor-repressor than Nrf1α, albeit both isoforms are endowed with an intrinsic capability to prevent tumor development and malignant growth.

### 3.8. Nrf1α and TCF11 Regulate Critical Genes for Improving the Survival Rate of HCC Patients

To weigh the practical effects of TCF11, Nrf1α and Nrf2 on human hepatocellular carcinoma (HCC), their relevancies to the overall survival (OS) of HCC patients were firstly investigated (Figure 8). For this, HCC-relevant molecules were selected from the UniProt database and their genes were further parsed by the Kaplan-Meier Plotter [55], along with other relevant databases [56–58], to find those markers of being significantly correlated with OS of patients with HCC. As shown in Figure 8 (*A1-A8*) and Table S10, a better OS was predicted to couple with increased expression of *CHP2* (*calcineurin like EF-hand protein 2*), *CPS1* (*carbamoyl-phosphate synthase 1*), *FOXO1* (*forkhead box O1*), *IRS4* (*insulin receptor substrate 4*) and/or *NKX2-8* (*NK2 homeobox 8*); this was also accompanied by reduced expression of *AKR1B10* (*aldo-keto reductase family 1 member B10*), *EPO* (*erythropoietin*), *MUTYH* (*mutY DNA glycosylase*) and/or *PKM* (*pyruvate kinase M1/*2). Besides, GP73 (also called GOLM1, golgi membrane protein 1) has been reported to be a potential diagnostic marker for primary HCC [59,60], while GPC3 (glypican 3) acts as another key biomarker for early diagnosis of human HCC and a rational immunotherapeutic target for HCC [61,62]. Although no significant differences in the expression levels of *Nrf1* or *Nrf2* amongst various tumor tissues from numerous patients with distinct stages of HCC were examined (Table S10), such two CNC-bZIP factors had been determined to differentially regulate the progression of HCC [17]. Notably, basal expression of Nrf1 in distinct HCC tissues had been shown significantly altered, relative to the equivalent expression in their adjacent para-carcinoma tissues or those expressed in normal liver cells [16]. These suggest that Nrf1 and Nrf2 execute distinct functions in the progression of HCC through differentially regulating putative pathophysiological processes. However, discrete isoforms of Nrf1 were also undistinguished in the above-described databases, but their unique activities are required for being further investigated separately.

**Figure 8.**
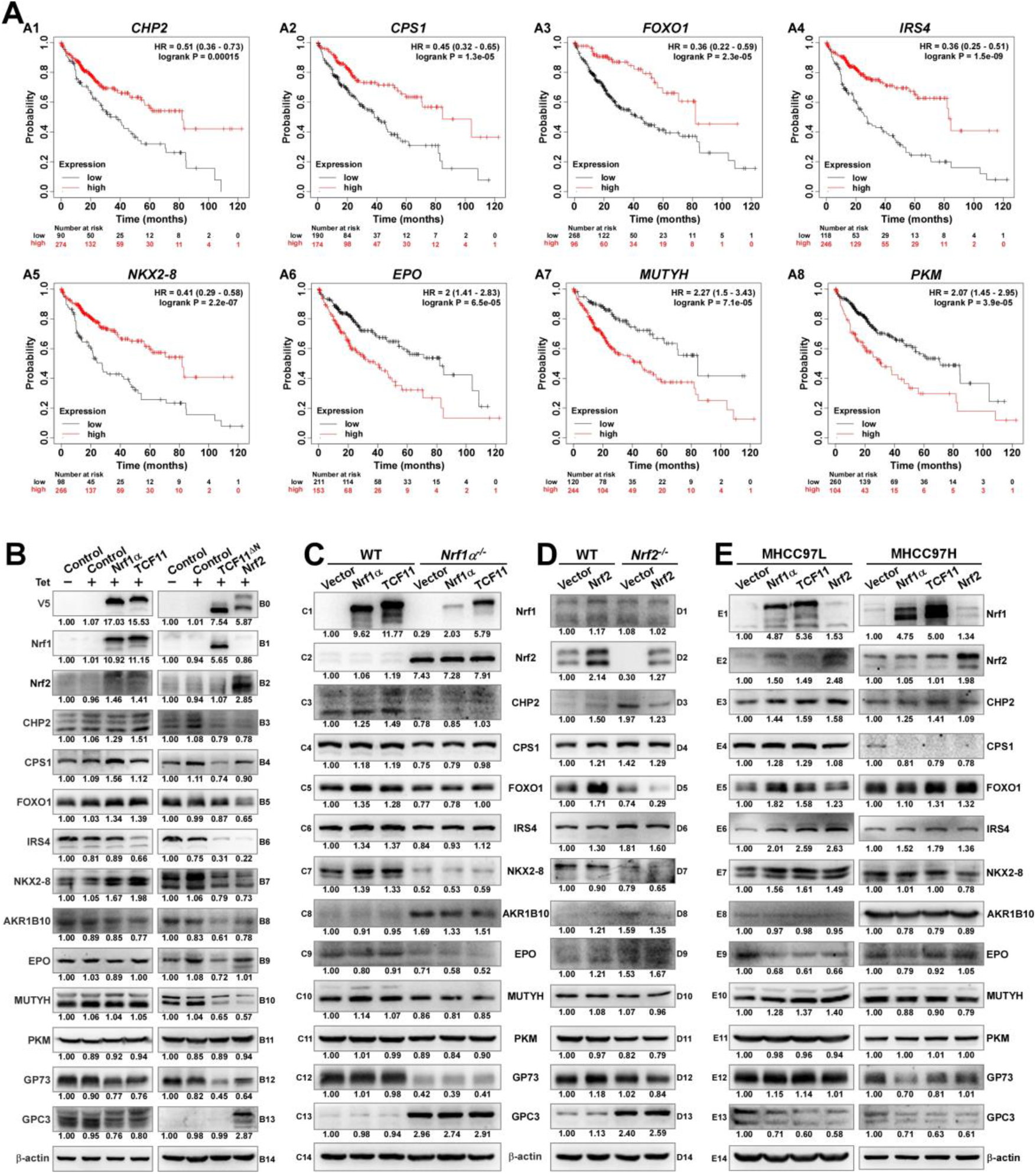
Diverse effects of Nrf1 and Nrf2 on those HCC-relevant proteins that are significantly correlated with the overall survival (OS). (**A**) The correlation analysis between of hepatocellular carcinoma (HCC) associated genes and the relevant OS rates of patients with HCC. (**B**) The protein expression levels of HCC-associated proteins in *Nrf1α*, *TCF11*, *TCF11^ΔN^* and *Nrf2* cell models were examined by Western blotting analysis of these experimental cell lines that had been induced with or without 1 μg/mL Tet for 12 h before being harvested. (**C**) Both wild type HepG2 and its derived *Nrf1α*^−/−^ cells were transfected with either Nrf1α or TCF11 expression plasmids, and then allowed for 24-recovery from transfection in the fresh medium before being subjected to Western blotting, to identify the protein levels of HCC-associated proteins. (**D**) Both wild type HepG2 and its derived *Nrf2*^−/−^ cells were transfected with an Nrf2 expression plasmid, and then allowed for 24-recovery from transfection in the fresh medium before being subjected to Western blotting. (**E**) MHCC97L or MHCC97H cells were transfected with each of Nrf1α, TCF11 and Nrf2 expression constructs, and then allowed for 24-recovery from transfection in the fresh medium before being examined by Western blotting to identify abundances of HCC-associated proteins as described above. The intensity of all the immunoblots was calculated and shown on the bottom (B to E).

The immunoblotting results showed that TCF11 and Nrf1α enabled the normal cells to increase CHP2 and CPS1 abundances, respectively (Figure 8, *B3* & *B4*). Although FOXO1 was slightly increased, NKX2-8 was markedly increased, by both TCF11 and Nrf1α (*B5* & *B7*), but IRS4 was down-regulated by TCF11 but not Nrf1α (*B6*). Furtherly, AKR1B10 and EPO were, to greater or less extents, diminished by TCF11 and Nrf1α (*B8* & *B9*). Notably, significant decreases in the two HCC biomarkers GP73 and GPC3 were caused by TCF11 and Nrf1α (*B12* & *B13*). By sharp contrast, most of the above-examined proteins were reduced by TCF11^ΔN^ and Nrf2 (Figure 8B, *right panels*), with an exception that GPC3 was significantly up-regulated by Nrf2, but not TCF11^ΔN^.

Further examinations revealed that overexpression of TCF11 and Nrf1α in HepG2 cells enabled CHP2, CPS1, FOXO1, IRS4 and NKX2-8 to be enhanced to varying extents, but they were markedly suppressed by loss of *Nrf1α^−/−^* (Figure 8C, *left panels*). Such loss of *Nrf1α* also gave rise to constructive enhancement of Nrf2, AKR1B10 and GPC3, as accompanied by constructive abolishment of EPO and GP73, besides CHP2 and NKX2-8. However, it is intriguing to note that all these constructive changes could not be ameliorated by modest restoration of ectopic TCF11 or Nrf1α into *Nrf1α^−/−^* cells (Figure 8C, *right panels*). By contrast, forced expression of Nrf2 in HepG2 cells caused only a marginal increase in FOXO1 or GP73, but the other examined genes were unaffected (Figure 8D, *left panels*). Conversely, loss of *Nrf2* caused a slight increase in CHP2 and GPC3; but the former CHP2 was reduced by ectopic Nrf2 restoration to its basal levels, while the latter GPC3 was not mitigated by ectopic Nrf2 (Figure 8D, *right panels*).

The above-described data indicate distinct contributions of TCF11, Nrf1α and Nrf2 to differential or opposite regulation of endogenous expression levels of different, even the same, target genes. This notion was herein substantiated by our further comparative experiments of TCF11, Nrf1α and Nrf2 that had been transfected for their respective overexpression in either MHCC97L or MHCC97H cell lines (Figure 8E). As anticipated, abundances of CHP2, CPS1, FOXO1, IRS4, NKX2-8 proteins were increased with their respectively-varying trends by ectopic TCF11, Nrf1α or Nrf2 expression in MHCC97L cells (Figure 8E, *left panels*), whereas such events did not occur in MHCC97H cells (Figure 8E, *right panels*). Besides, both EPO and GPC3 were suppressed by ectopic expression of TCF11, Nrf1α and Nrf2 in MHCC97L cells, but almost unaffected in MHCC97H cells with hyper-expression of Nrf2. Such discrepancies in these gene expression are much likely contributable to distinct metastatic potentials and other malignant properties of between MHCC97L and MHCC97H cell lines. Subsequently, an integrated interaction network regulated by Nrf1 and Nrf2 (Figure 9A) was built on the basis of the protein interaction database STRING [63], in combination with our experimental results as shown above. The distinct expression levels of those indicated genes that make up the network in stable expression cells (Figure 9B) or knockout model cells (Figure 9C), were also indicated in the heatmap, respectively. Taken altogether, it is demonstrated that Nrf1α and TCF11, but not Nrf2, are conferred for an intrinsic capability to regulate those genes critical for improving the survival rate of patients with HCC, such that Nrf1α or TCF11 perform a strikingly disparate effect from that of Nrf2 on human hepatoma (Figure 9D).

**Figure 9.**
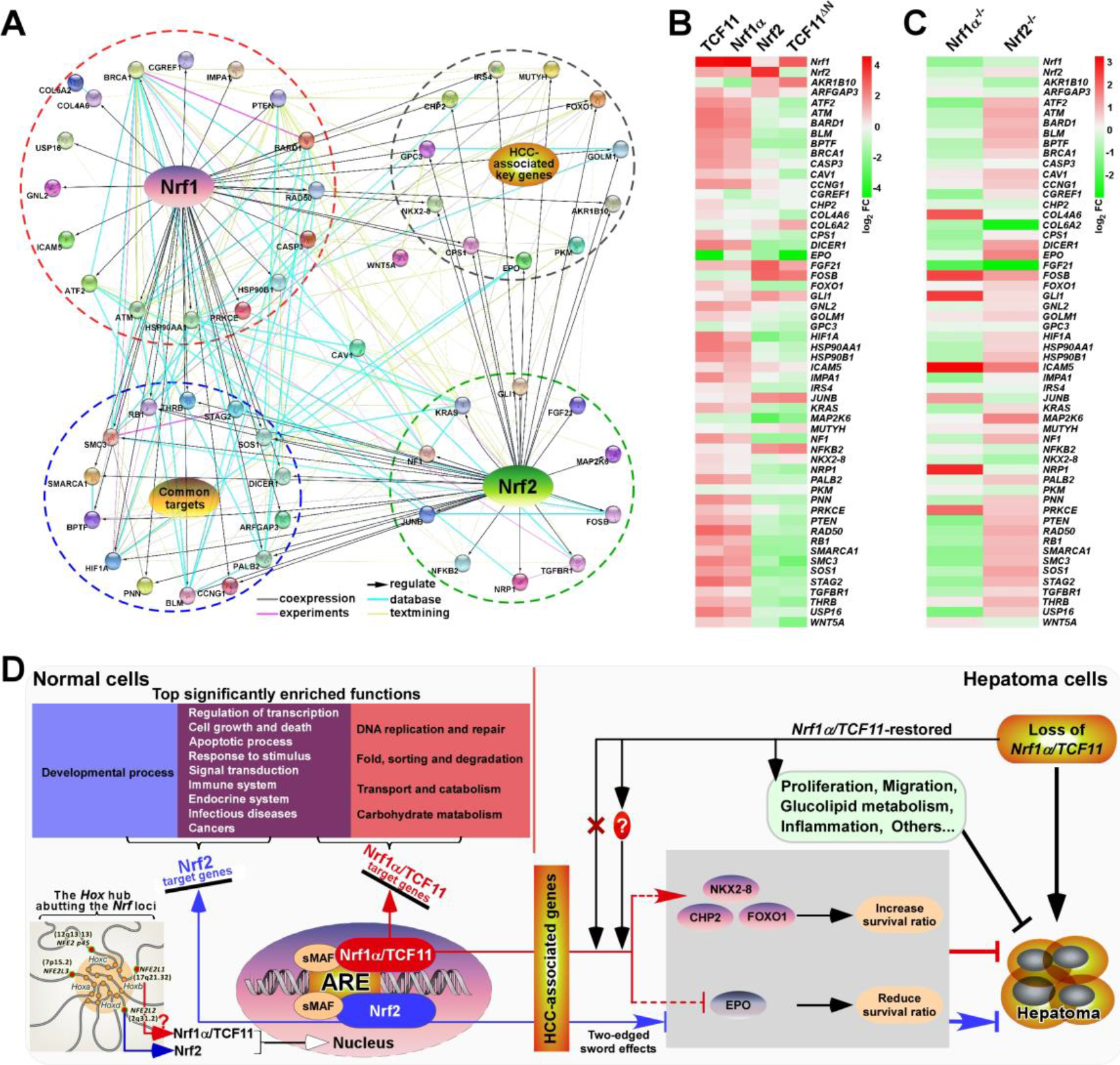
Relationship and difference between the regulation patterns of Nrf1 and Nrf2 on their targets. (**A**) The functional protein association networks of targets regulated by Nrf1 or Nrf2. Of note, the protein-protein associations are determined by various ways, which are thus represented by different colored edges as indicated. (**B**) The heatmap of the sequencing expression of genes, which are composed of the network, with distinct expression levels in *TCF11*, *Nrf1α*, *Nrf2* and *TCF11^ΔN^* cell lines. The color of the nodes in the heatmap represents the value of log_2_ (fold change) as shown in the color bars, indicating the gene expression trend as compared with the control group (upregulation or downregulation, were marked in red or green, respectively). (**C**) The heatmap of the sequencing expression of genes in *Nrf1α^−/−^* and *Nrf2^−/−^* cell lines. The color of the nodes in the heatmap represents the value of log_2_ (fold change) as shown in the color bars, indicating the gene expression trend as compared with the wild-type HepG2 cells. (**D**) A comprehensive regulatory model is proposed to reveal the different effects of Nrf1 and Nrf2 on hepatoma (*right panel*). In addition, the *Hox* hub abutting the distinct *Nrf* loci was also indicated (*at the lower left corner*).

## 4. Discussion

In this study, we have established four controllable cell models for stably expressing *TCF11*, *TCF11^ΔN^*, *Nrf1α* or *Nrf2*, which all occur naturally with their respective intact structural domains, albeit they were tagged C-terminally by a neutral V5 epitope. Of striking note, Nrf1α and TCF11 are two longer isoforms of the ER-resident Nrf1 with changing gears in their overall transcriptional activity, because both proteins are tightly controlled by their unique topobiological rheostat modules from the ER to enter the nucleus. For instance, the NST (Asn/Ser/Thr-rich) domain transactivation of Nrf1α/TCF11, as well as SKN-1 (Skinhead-1) and CncC (Cap‘n’collar isoform C), is also required for re-editing of their indicated asparagines to aspartates in this region during dynamic dislocation from the lumen to extra-ER subcellular compartments [64,65]. Upon lack of such amino acid re-editing by peptide: N-glycosidase (PNG1), this causes an evident decrease in the rheostat capacity of Nrf1α/TCF11, and even loss of *PNG1* or its mutations results in inactivation of Nrf1α/TCF11, manifesting inflammation and adrenal insufficiency [66–68]. Furtherly, the rheostat capacity of Nrf1α/TCF11 is also monitored by its reversible ubiquitination and deubiquitination during its selective ER-associated proteasome-regulated processing to multiple isoforms [69,70].

During our manuscript preparation, removal of the N-terminal ER signal peptide-containing 104-aa or 121-aa regions from Nrf1α/TCF11 to yield two artificially-truncated mutants (i.e., Nrf1^ΔN104^ or Nrf1^ΔN121^) had been reported by both independent Bollong’s and Ooi’s laboratories [44,45]. It is, to our great surprise, seen that such artificial mutants Nrf1^ΔN104^ or Nrf1^ΔN121^ were asserted to represent two constitutive processed Nrf1 activators, although they each retain a negative PEST region. Thereby, our study of *TCF11^ΔN^* showed an N-terminal 2-156 residues-truncated mutant (i.e., TCF11^ΔN2-156^), which may also arise from a naturally-splicing transcript [43]. However, TCF11^ΔN^ is herein identified as a mimic Nrf2 factor, because both share conserved structural domains (Figure 1A). This seems to totally contradict the purpose of Nrf1^ΔN104^ or Nrf1^ΔN121^ by both Bollong’s and Ooi’s colleagues [44,45]. Additional two mutants of caNrf2 to block its Keap1-binding activity were also utilized in their experimental settings. Besides, it should be of crucial importance to note that a strong acidic property of 3×Flag, wherever it was tagged at the N-terminal or C-terminal ends of a given protein, exerts distinctive or even opposing effects on its topobiological folding and dynamic moving in and out of membranes and other subcellular compartments [69,71]. Overall, our study has provided four cell tools to evaluate similarities yet differences in between intact TCF11, TCF11^ΔN^, Nrf1α and Nrf2-regulatory profiles, which are stimulated at the Tet-inducibly controllable levels.

Subsequently, these four cell lines were determined by transcriptomic sequencing in an integrative combination with routine reductionist approaches. As expected, comprehensive functional annotation of TCF11, TCF11^ΔN^, Nrf1α and Nrf2 unraveled that they are likely to perform diverse biological functions by combinationally overlapping or competitively opposing modules within distinct cellular networks, apart from their unique functions. Thereby, a further understanding of their respectively regulated targets and functions should be achieved only by making one of them function alone. For this end, our established cell lines stably expressing *TCF11*, *TCF11^ΔN^*, *Nrf1α* or *Nrf2* respectively, are an invaluable tool to gain insights into distinguishing their regulatory profiles, while they are allowed to function alone. Consequently, the regulatory patterns of Nrf1α and TCF11 are similar yet different, which are strikingly disparate from that of Nrf2, even with certain opposite effects on the same targets. Such distinctions between Nrf1α/TCF11 and Nrf2 are attributable to differences in their primary structures and functional subcellular locations. This is due to the objective fact that the ER-associated domains of Nrf1α/TCF11 determine its unique membrane topobiology and post-synthetic processing mechanisms, which enables it to be distinguishable from Nrf2. Yet, once the ER-targeting signal peptide-adjoining region were deleted to yield TCF11^ΔN^, this N-terminally truncated isoform does exhibit a similar regulatory pattern to that of Nrf2. However, this notion appears to totally contradict the conclusions drawn from two so-called constitutive activators Nrf1^ΔN104^ or Nrf1^ΔN121^ by Bollong’s and Ooi’s groups [44,45]. As such, their analyses by combining ChIP-seq and RNA-seq data revealed that Nrf1^ΔN121^ (or Nrf1^ΔN104^) had 3.19-fold numbers of target-binding peaks than those of caNrf2-binding targets. Of note, over 75% of Nrf1^ΔN121^-binding targets are focused preferentially on the consensus ARE sequence (5’-TGAC/GnnnGC-3’) flanked by AT-enriched motifs, while only less than 45% of caNrf2-binding targets are recruited on ARE-like sequences flanked by GC-enriched motifs [44]. It is inferable that Nrf1 exerts its unique biological functions by predominantly regulating ARE-driven cognate genes, whereas Nrf2 has a widely-varying capacity to elicit its promiscuous roles in diversely mediating non-ARE-battery genes. Altogether, with our previous work [17,37] demonstrating that both Nrf1α/TCF11 and Nrf2 can mutually influence each other by their inter-regulating at distinct levels, this leads to the formation of a steady-state regulatory network system finely monitored by a negative feedback loop to maintain the cell homeostasis and organ integrity.

Since early discovery of Nrf2 required for regulating antioxidant and detoxification genes [72], the overwhelming majority of investigations in this field have been focused disproportionately on this CNC-bZIP factor and its negative regulator Keap1 in response to redox stress, but also provided myriad insights into its promiscuous roles in biology and medicine, as well as drug development [73–75]. However, aside from these studies by routine reductionist approaches, the whole genome-widely integrative analyses uncovered the underlying facts that Nrf1, but not Nrf2, is essentially required for regulating important homeostatic and metabolic genes involved in the normal growth and development throughout life process. This is fully consistent with the experimental evidence showing that loss of Nrf1 results in mouse embryonic lethality and also causes adult pathological phenotypes varying within its gene-targeting mutant organs (reviewed by [70]). Such bad consequences are attributable to severe endogenous oxidative stress and fatal defects in redox metabolism reprogramming and relevant constitutive gene expression profiles of *Nrf1*-deficient cells (this study and [12]). Collectively, these facts demonstrate that Nrf1 is a predominant determinist factor of cellular constitutive redox metabolic homeostasis with organ integrity. That is to say, Nrf1 has a potent capacity to contribute to the steady-state robustness of cell physiological homeostasis. By contrast, Nrf2 is much likely to make a major contribution to the sensitive plasticity of cell homeostasis, because it has been accepted as a master regulator in response to diverse stresses. This is also supported by the facts of no obvious phenotypes resulting from global loss of *Nrf2* in mice, but its deficiency leads to more susceptibility to chemical carcinogens and diverse stresses than wild-type controls [76,77].

Such differential contributions of Nrf1 and Nrf2 to cell homeostasis robustness and plasticity are also likely resulted from a striking evolutionary conservation of their CNC-bZIP family in distinct species during nature selection. Comparative genomics analyses revealed that, although this CNC-bZIP family expansion and diversification have occurred in vertebrates, Nrf1 is viewed as a living fossil that is more ancient than Nrf2, because it is conserved closely to the only CNC in *Drosophila melanogaster*, SKN-1 in *Caenorhabditis elegans* and each of newly-identified Nach factors in simple multicellular eukaryotes, but not in unicellular protozoans or other prokaryotes beyond *Endozoicomonas* [4]. This implies that they have evolved for multicellular cooperative selection. For instance, the *C. elegans* SKN-1, like Nrf1, is selectively processed by post-transcriptional and post-translational ways to yield distinct isoforms, one of which is N-terminally truncated (by analogy of TCF11^ΔN^) but manifested the Nrf2-like functions, albeit with no existence of Keap1-like orthologues [65,78]. Such the highly evolutional conservativity of Nrf1, but not of Nrf2, strongly demonstrates that it is indispensable for basal constitutive contributions to orchestrating the ensemble of critical gene regulatory networks, such that the normal cell homeostasis and organ integrity have been perpetuated in the entire course of life.

In fact, the CNC-bZIP family members (p45, Nrf1, Nrf2 and Nrf3) are, though diversified in vertebrates, topologically organized together with the developmental *Hox* gene clusters (as shown in Figure 9D), upon forming the PcG hub to maintain their genes in a silent-but-poised state [79]. A recent ChIP data analysis revealed that most of Nrf1-binding sites are focused closely to the canonical ARE sequences that are widely located in its cognate gene promoter and distal intergenic enhancer regions (i.e., dynamic genome-topology as a functional primer is built by spatial interactions of distal enhancers with the proximal promoters in the same genes or even between different genes in chromosomal architecture), whereas Nrf2-binding sites are promiscuously loosed to ARE-like or no-ARE sequences that are, however, constrained narrowly in its target gene promoters [44]. Collectively, the interplay between selective transcription factor (e.g., CNC-bZIP)-regulated gene programming and genome conformation is surmised as a driving force for cell-fate decision with distinct type-featured identifications.

Notably, the versatile Nrf2 acts *de facto* as a promiscuous, not essential, player in its biology, because it is dispensable for normal growth and development in mice with no phenotypes (e.g., cancer) resulting from its genetic loss [74,77]. Contrarily, accumulating evidence clearly demonstrates that hyperactive Nrf2 promoted cancer development and malignance, because it is relevant to most of cancer hallmarks [73]. This implies that, except that Nrf2 can exert a significant cytoprotective function against diverse stresses so as to confer the cells to be acquired for the adaptive responses, its long-term hyper-activation can also critically inspire potential cancerous cells to be extricated from being rigidly controlled and confined by inevitability of the host multicellular cooperative evolution. The notion is supported by transcriptome sequencing of *Nrf2*-deficient cells, because this loss of Nrf2 causes hepatoma to be significantly ameliorated or completely prevented [17]. In the other way round, aberrant accumulation of hyperactive Nrf2 in *Nrf1*-deficient cells (or livers) leads to further malignant transformation of human hepatocellular carcinoma (HCC).

In particular, the spontaneous development of NASH-based inflammation and hepatoma are resulted from severe endogenous oxidative stress and fatal defects in basal constitutive gene expression affected by its genetic instability in mouse *Nrf1*-deficient livers [80–82]. Of note, TCF11 is absent in mice and has a low proportion in all human hepatoma cell lines, but it, together with Nrf1α at a 1:1 ratio, exists in human normal cells (Figure S9). Upon *Nrf1α*-specific knockout from hepatoma cells, this results in cancer malignant growth and metastasis to the lungs in xenograft model mice [16,54]. The *Nrf1α^−/−^*-derived cancer deterioration is definitely resulted from severe oxidative stress, fateful defects in the redox metabolism reprogramming, and marked dysfunctions of fate-decisive gene regulatory networks, along with critical aberrant signaling transduction networks [12,17]. However, it is, to our great surprise, found that *Nrf1α^−/−^*-led malignant transformation is markedly alleviated and prevented by silencing of Nrf2; this is also accompanied by blocking oxidative stress upon Nrf2 knockdown [12,17]. More excitingly, restoration of *Nrf1α* is allowed for significant mitigation of *Nrf1α^−/−^*-exacerbated tumor to similar wild-type extents, while its malignant growth is further suppressed or even abolished by *TCF11* restoration, with complete coagulative necrosis of tumor tissues (Figures 7B & S8). This demonstrates that TCF11 is a potent tumor-suppressor than Nrf1α, while Nrf2 is a tumor-promotor, particularly in the *Nrf1α^−/−^* case. Altogether, loss of Nrf1α/TCF11’s function, with hyperactive Nrf2 accumulation, results in liver cancer initiation and progression. This consequence is likely originated from endogenous oxidative stress-induced damages of mutant cells in aberrant metabolic inflammatory microenvironments. This gives rise to an evolvable selection force in so much of *Darwinian* dynamics, so that a clade of the mutant cancerous-prone cells is endowed with a self-defined fitness function and also specified to acquire for an independent of the host team optimum cooperative confinements, in order to behave themselves with own properties of cell division, proliferation and migration during carcinogenesis and progression.

Moreover, this work further demonstrates that the tumor-preventing effect of Nrf1α and TCF11 is accompanied by the constitutive activation or repression of critical genes for improving the overall survival of patients with hepatocellular carcinoma (Figure 9). This is because these changes can be ameliorated by either Nrf1α- or TCF11-restored lines (as mentioned in Figure 7). However, their activation or repression of some genes could not be ameliorated by compensation of ectopic *Nrf1α* and *TCF11* transfection into those *Nrf1α/TCF11*-deficient cancer cells. This is surmised to be the relevance of those given genes closely to the contexts within genome topological conformations, enabling ectopic Nrf1α/TCF11 factors to be allowed or forbidden for direct access to target genes (as shown in the lower left corner of Figure 9D).

Lastly, it should also be noted that our subcellular fractionation and immunofluorescence results have unraveled that a certain fraction of TCF11, TCF11^ΔN^, Nrf1α or Nrf2 can be allowed for spatial translocation from the cytoplasmic to the nuclear compartments, in which they gain access to target genes (Figure S3). Interestingly, these nuclearly-located fractions of TCF11, TCF11^ΔN^, Nrf1α or Nrf2 are rapidly degraded. This is due to this fact that inhibition of their proteasomal degradation by MG132 causes an obvious increase in each protein expression level of TCF11, TCF11^ΔN^, Nrf1α or Nrf2, and they were also markedly recovered in their nuclear fractions of MG132-treated cells (Figure S3). Altogether, these demonstrate that only after TCF11, TCF11^ΔN^, Nrf1α or Nrf2 enter the nucleus and stay for a given time in this subcellular compartment, they are conferred to exert their putative physiological functions by regulating transcriptional expression profiles of distinct subsets of target genes. On the contrary, it is inferable that if they are degraded totally, as reported previously, in the extra-nuclear cytoplasmic compartments, just under normal physiological conditions, none of their remaining proteins can be allowed for dynamic repositioning into the nucleus, so that no physiological functions of cognate genes would be regulated by these CNC-bZIP factors. However, this notion cannot hold true in fact, albeit the detailed mechanisms are required to be further explored.

## 5. Conclusions

In summary, four useful model cell lines stably expressing *TCF11*, *TCF11^ΔN^*, *Nrf1α* or *Nrf2*, not mutants, are yielded herein. Their transcriptional expression levels are controlled by a tetracycline-inducible switch, but not monitored by one of redox inducers (e.g., sulforaphane or *tert*-Butylhydroquinone) or ER stressors (e.g., tunicamycin or thapsigargin). These cells were subjected to the integrative systems biology analyses of their omics data with routine reductionist approaches. To explore the essential distinctions between Nrf1α/TCF11 and Nrf2 in their contributions to critical constitutive gene expression profiles for basal redox metabolism, normal growth, development, cell homeostasis and organ integrity, we have provided holistic relevant data as much as possible, in the present digital network era. Notably, some seemingly-paradoxical data are still retained, because they may serve as vital nodes of a few negative feedback (and feedforward) regulatory circuits existing in the self-organizing systems of life. This is for the sake of an objective truth in life. Significantly, it is demonstrated that TCF11 serves as a more potent tumor-suppressor than Nrf1α at preventing cancer development and progression. This is defined by similar yet different regulatory profiles of both isoforms, with a striking disparity from Nrf2. Rather, a naturally-spliced mutant TCF11^ΔN^ resembles Nrf2 with largely consistent structure and function in regulating similar sets of target genes. Interestingly, the tumor-preventing effect of Nrf1α/TCF11 seems to be accompanied by certain constitutive activation or repression of critical genes for improving the overall survival rates of patients with hepatoma. Once loss of Nrf1α/TCF11’s function, with hyperactive Nrf2 accumulated, this leads to severe endogenous oxidative damages, aberrant redox metabolic inflammation, and ultimate spontaneous hepatoma. Such genetic and nongenetic drivers could be integrated as a selection force in *Darwinian* dynamics to enable for stochastic speciation of *Nrf1*-deficient cells during carcinogenesis and ensuing progression, albeit this is required for deeply studies. Taken together, this study provides a holistic perspective to give a better understanding of essential differences between Nrf1α/TCF11 and Nrf2 in biology and medicine. Thereby, this facilitates drug discovery to induce Nrf1α/TCF11 as a new potent chemoprevention target against cancer.

## Supplementary Materials

Figure S1: Structural comparison of distinct Nrf1 isoforms with Nrf2. Figure S2: A detailed diagram of Flp-In system. Figure S3: The distribution of *TCF11*, *TCF11^ΔN^*, *Nrf1α* and *Nrf2* in each of their stably-expressing cells. Figure S4: Statistical analysis of the transcriptome sequencing data. Figure S5: Identification of a significant correlation between real-time quantitative PCR results and transcriptome sequencing data. Figure S6: Similarities and differences amongst those mainly regulatory profiles of TCF11, TCF11^ΔN^, Nrf1α and Nrf2. Figure S7: Alterations in the cell cycle and apoptosis caused by restoration of *Nrf1α* and *TCF11*. Figure S8: The HE staining results of tumors in each group. Figure S9: The relative proportion of *Nrf1α* and *TCF11* in different liver cancer cells and normal cells. Table S1: The primers for construction or qPCR used in this paper. Table S2: The differentially expressed genes regulated by TCF11. Table S3: The differentially expressed genes regulated by Nrf1α. Table S4: The differentially expressed genes regulated by Nrf2. Table S5: The differentially expressed genes regulated by TCF11^ΔN^. Table S6: The functional annotation of DEGs from group A to C. Table S7: The functional annotation of DEGs from group D to F. Table S8: The functional annotation of DEGs from group G to I. Table S9: The functional annotation of DEGs from group J to L. Table S10: Analysis of the significance of the potential HCC-associated proteins in relation of liver cancer in different databases.

## Author Contributions

M.W. designed and performed most of the experiments and bioinformatic analyses, made all figures and wrote this manuscript draft. Y.R. constructed the *Nrf1α*- and *TCF11*-restored cell lines and performed the subcutaneous tumor xenografts experiment. Both S.H. and K.L. helped M.W. with some molecular cloning to create several expression constructs and performed western blotting. L.Q. established both *Nrf1α^−/−^* and *Nrf2^−/−^* cell lines. Lastly, Y.Z. designed and supervised this study, analyzed and parsed all the data, helped to prepare all figures, wrote and revised this manuscript.

## Data Availability Statement

All data needed to evaluate the conclusions in the paper are present in this publication along with the supplementary information documents that can be found online. The additional data related to this paper may also be requested from the corresponding author on reasonable request. Besides, it should be noted that the preprinted versions of this paper had been initially posted at *bioRxiv* 2021.01.12.426360 (doi: https://doi.org/10.1101/2021.01.12.426360).

