## Supplemental Figures for "TCF11 Has a Potent Tumor-Repressing Effect than Its Prototypic Nrf1α by Definition of both Similar yet Different Regulatory Profiles, with a Striking Disparity from Nrf2"

Figure S1:

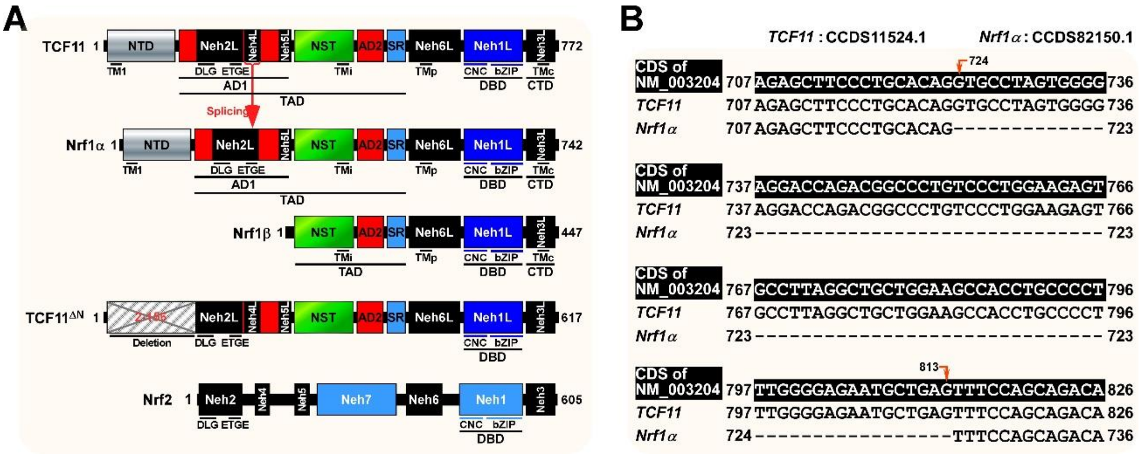


**Figure S1. Structural comparison among distinct Nrf1 isoforms and Nrf2.** (A) Schematics of the similarities and differences among structural domains of distinct Nrf1 isoforms and Nrf2. (B) Alignment among the cDNA sequences of *Nrf1α*, *TCF11* and NM_003204 to show the missing part of *Nrf1α*.

Figure S2:

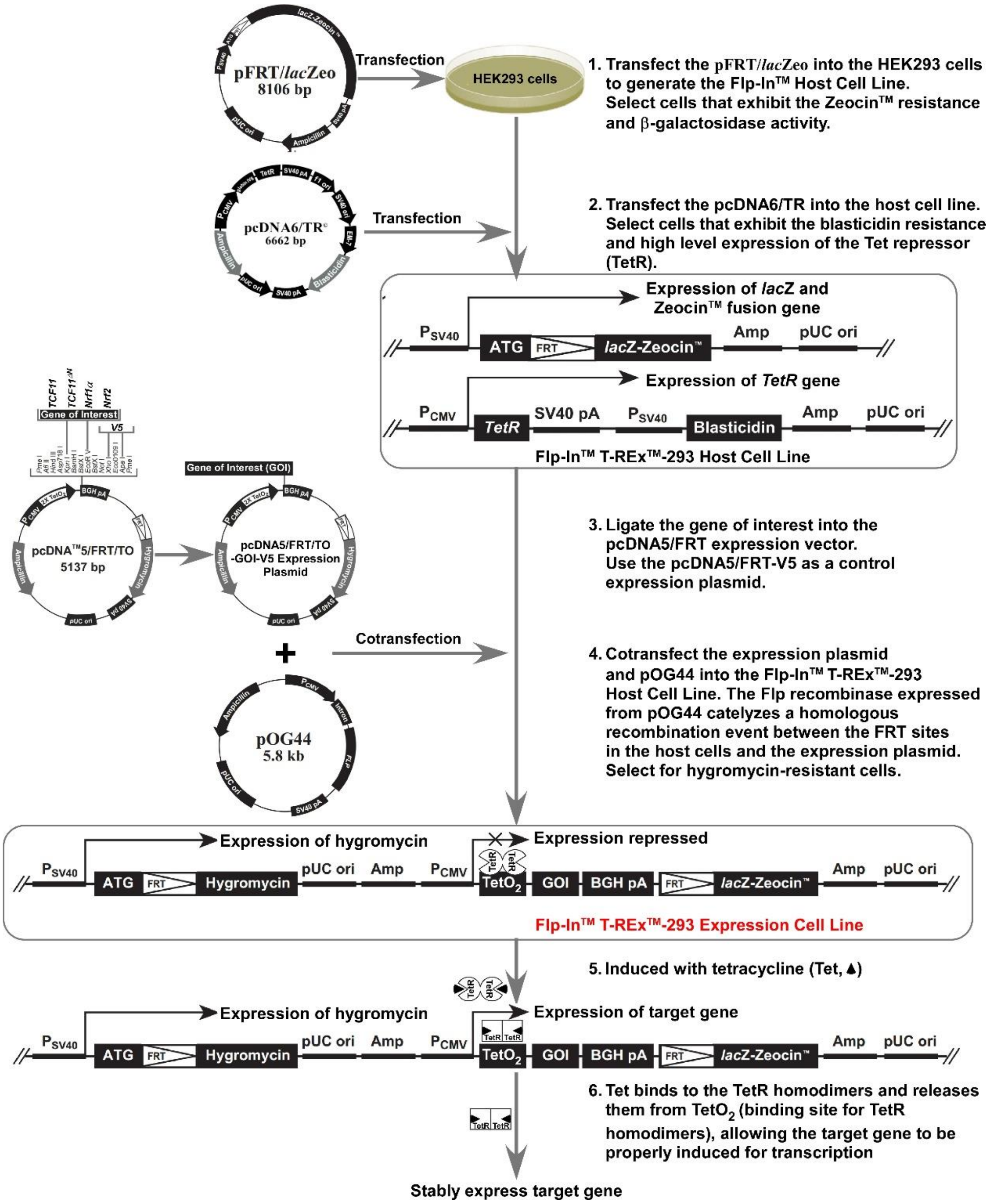


**Figure S2. Detailed diagram to establish the Flp-In™ T-REx™-293 expression cell lines with Flp-In system.** Firstly, transfect the pFRT/*lac*Zeo plasmid, which introduces a signal Flp Recombination Target (FRT) site into the genome and stably express the *lac*Z-Zeocin™ fusion gene under the control of the SV40 early promoter, into HEK293 cells, then select the Flp-In™ host cells that exhibit the Zeocin™ resistance and high β-galactosidase activity. Next, transfect the pCDNA6/TR plasmid, which stably expresses the Tet repressor gene (TetR) under the control of the constitutive human cytomegalovirus (CMV) immediate-early promoter, into the Flp-In™ host cells, and select the Flp-In™ T-REx™-293 host cells that exhibit the blasticidin resistance and high-level expression of TetR. After using the pCDNA™5/FRT/TO vector to construct the expression plasmid containing the gene of interest (GOI), the expression plasmid and pOG44 vector (which expresses the Flp recombinase under the control of the human CMV promoter) were co-transfected into Flp-In™ T-REx™-293 host cell line. The Flp recombinase expressed from pOG44 catalyzes a homologous recombination event between the FRT sites in the host cells and the pCDNA™5/FRT/TO expression plasmid, followed by the integration of the expression construct allows transcription of target gene and confers hygromycin resistance to the cells to generate the Flp-In™ T-REx™-293 expression cell line. Whilst the TetR will form homodimers and bind to Tet operator 2 ($TetO_2$) sequences in the pCDNA™5/FRT/TO expression plasmid to repress the transcription of target gene in expression cell line, only when the expression cells induced with tetracycline (Tet, binds to TetR homodimers causing a conformational change and release from the $TetO_2$ sequences) can the target gene be successfully transcribed and expressed stably.

### Figure S3:

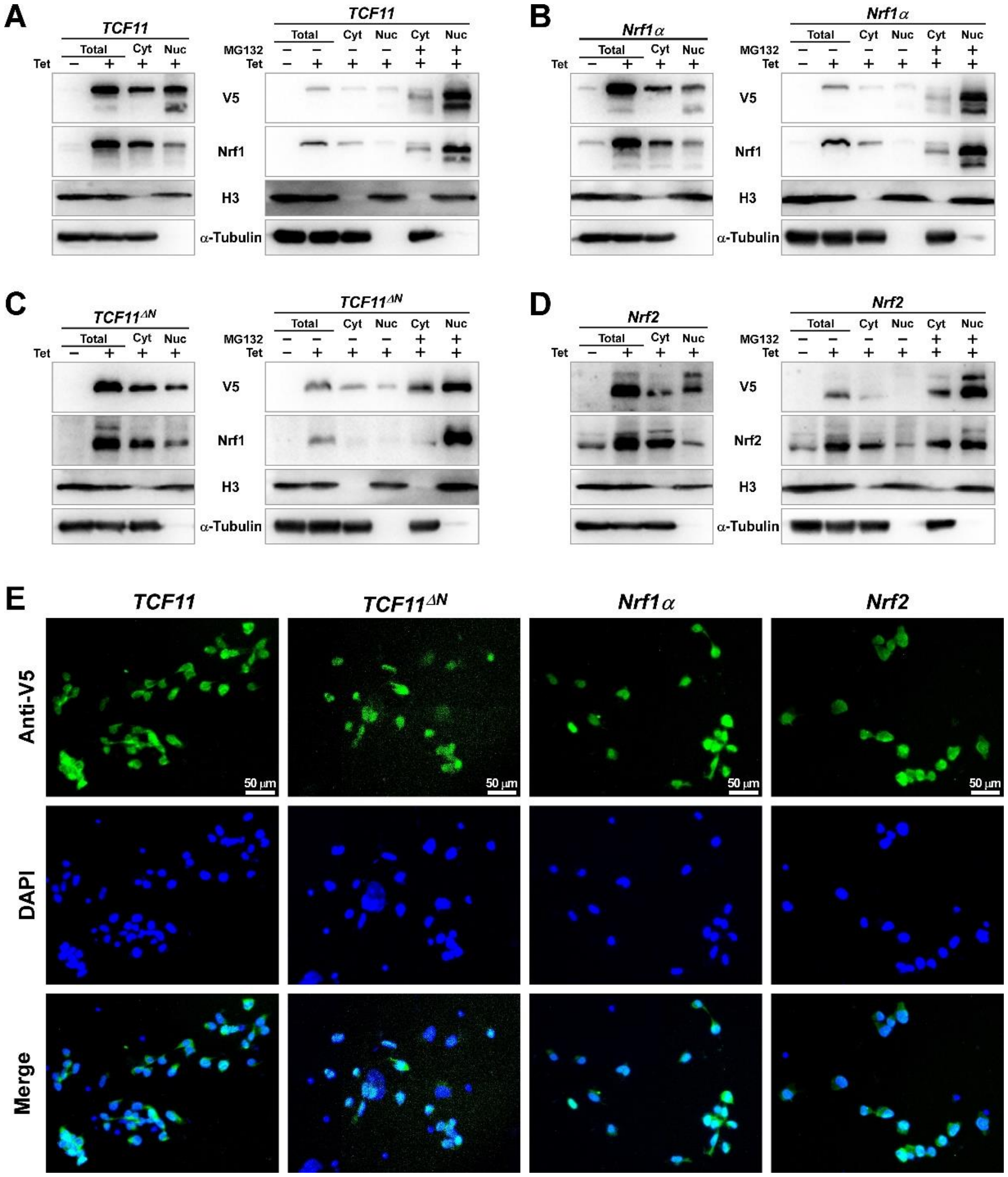


**Figure S3. The distribution of TCF11, TCF11$^{\Delta N}$, Nrf1α and Nrf2 in cells was detected by subcellular fractionation and immunofluorescence.** (A-D) After collecting the cytoplasmic and nuclear fractions of different cells that had been induced with Tet (1 μg/ml) for 12 h alone or in combination with MG132 (10 μmol/L, added in the last four hours of the 12 hours), Western blotting experiments were used to compare the expressions of TCF11 (A), Nrf1α (B), TCF11$^{\Delta N}$ (C) and Nrf2 (D) in the cytoplasm and nucleus. (E) Fluorescence images of TCF11, TCF11$^{\Delta N}$, Nrf1α and Nrf2 in distinct subcellular distributions were acquired by immunofluorescence with the primary antibody against V5, along with FITC-labeled second antibody. The nuclear DNA was stained by DAPI.

Figure S4:

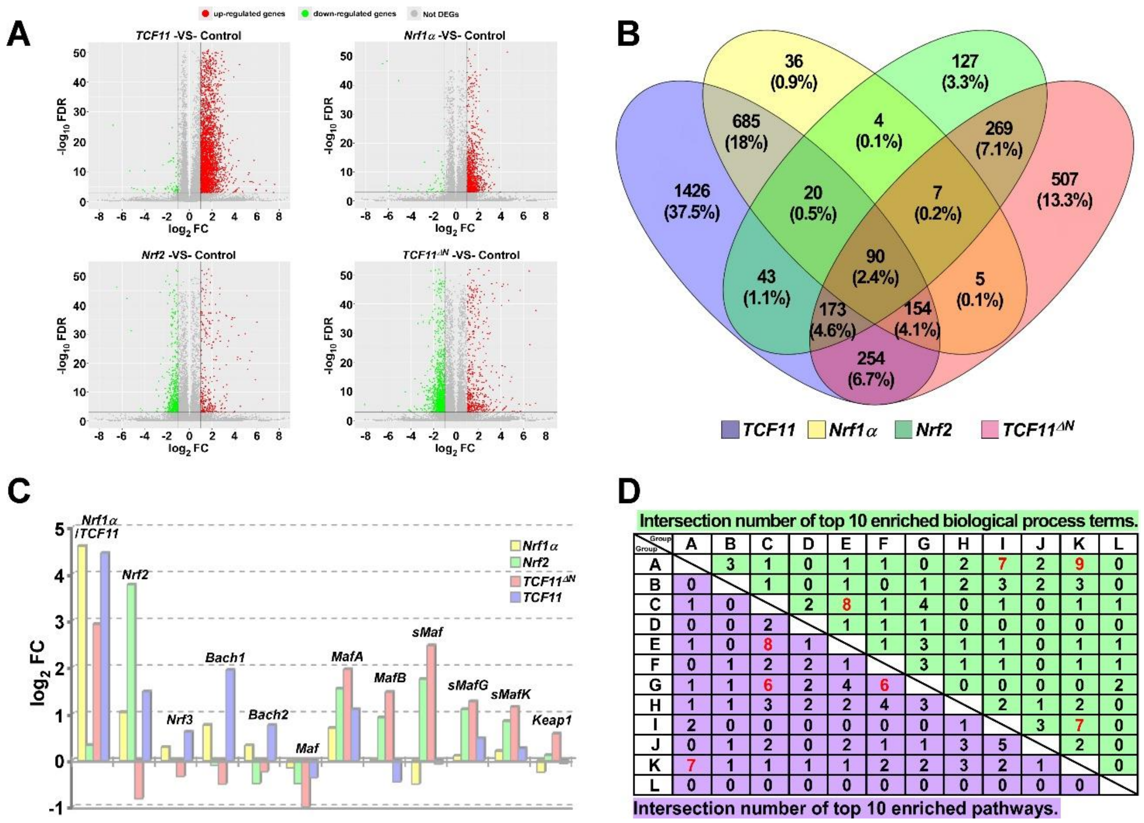


**Figure S4. The global statistical analysis of sequencing data.** (A) Scatter plots of global expression change trends of all genes detected in *TCF11*, *Nrf1α*, *Nrf2* and *TCF11*$^{\Delta N}$ cells. (B) The common or unique DEGs among sequenced samples were exhibited in the Venn chart. (C) The expression comparison of *Nrf1*, *Nrf2* and *Nrf3*, *Bach1* and *Bach2*, together with sMaf family members. (D) The number of common functions that enriched by DEGs in group A-L.

Figure S5:

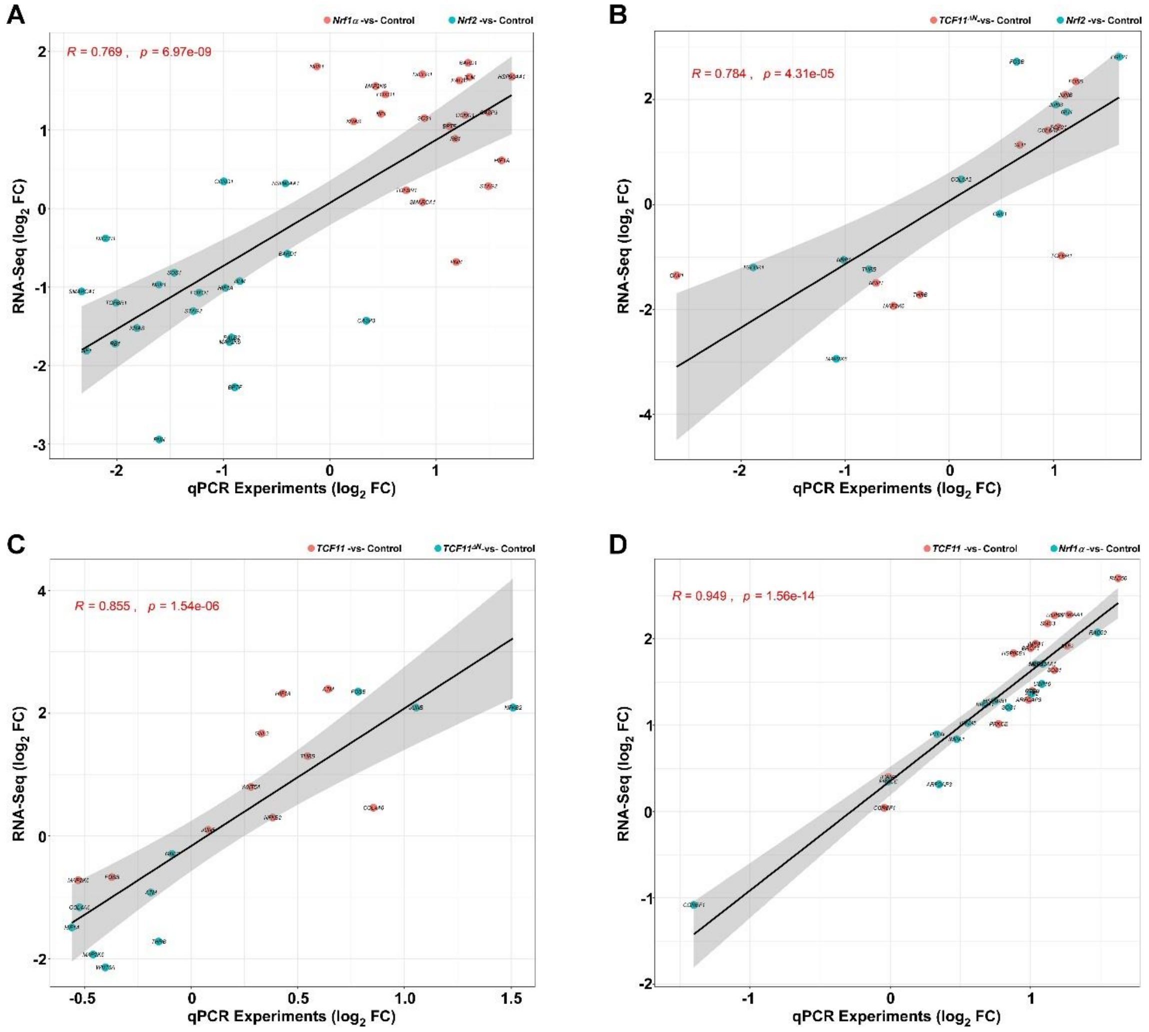


**Figure S5. Identification of correlation between real-time quantitative PCR results and transcriptome sequencing data.** The correlation analysis between the expression changes of genes in group A-C (A), D-F (B), G-I (C)and J-L (D) in qPCR experiments and transcriptome sequencing.

Figure S6:

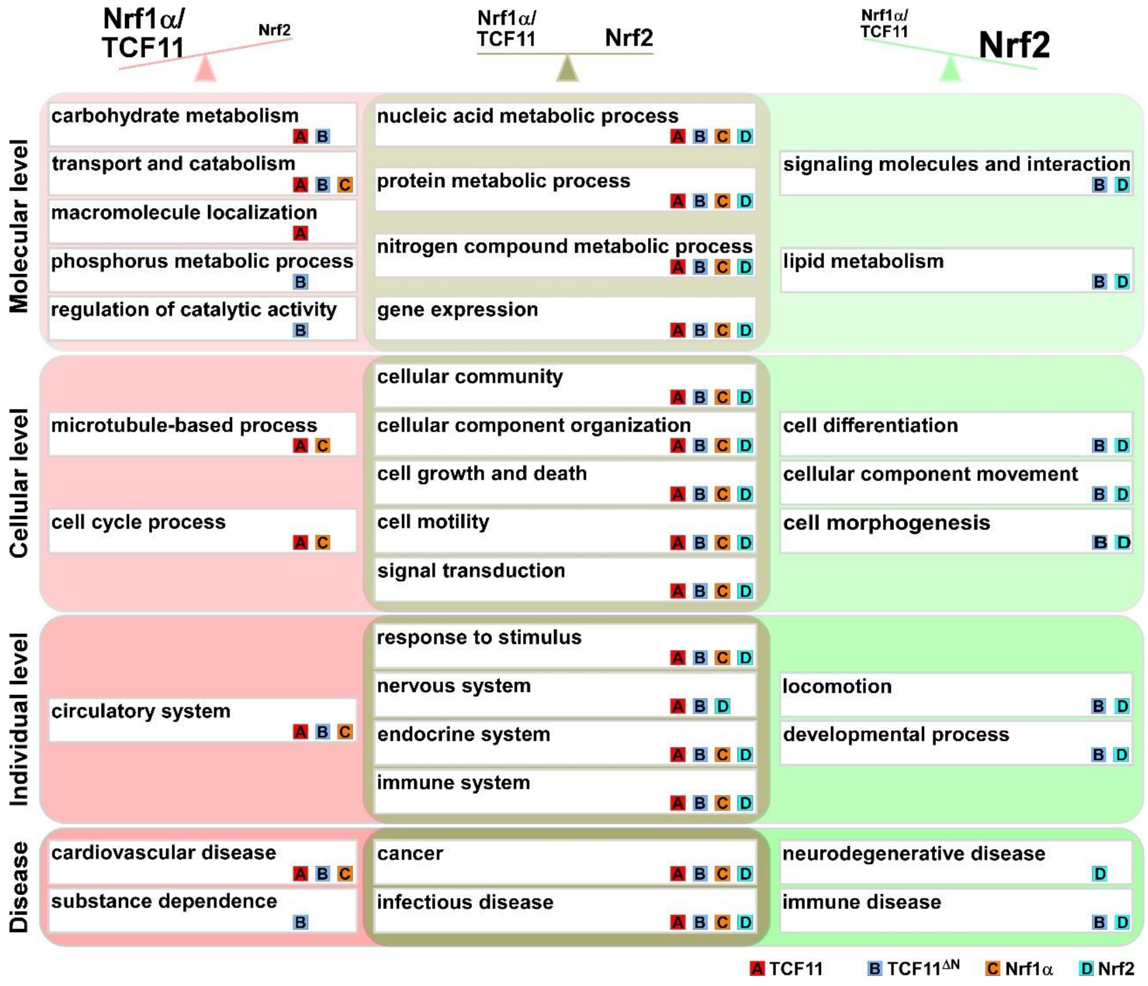


**Figure S6. The similarities and differences in the regulatory profiles of TCF11, TCF11$^{\Delta N}$, Nrf1α and Nrf2.** By summarizing the enrichment analysis of functional annotation of differentially expressed genes in each group, each transcription factor was labeled on the lower right side of the biological process that it can significantly participate in regulation.

Figure S7:

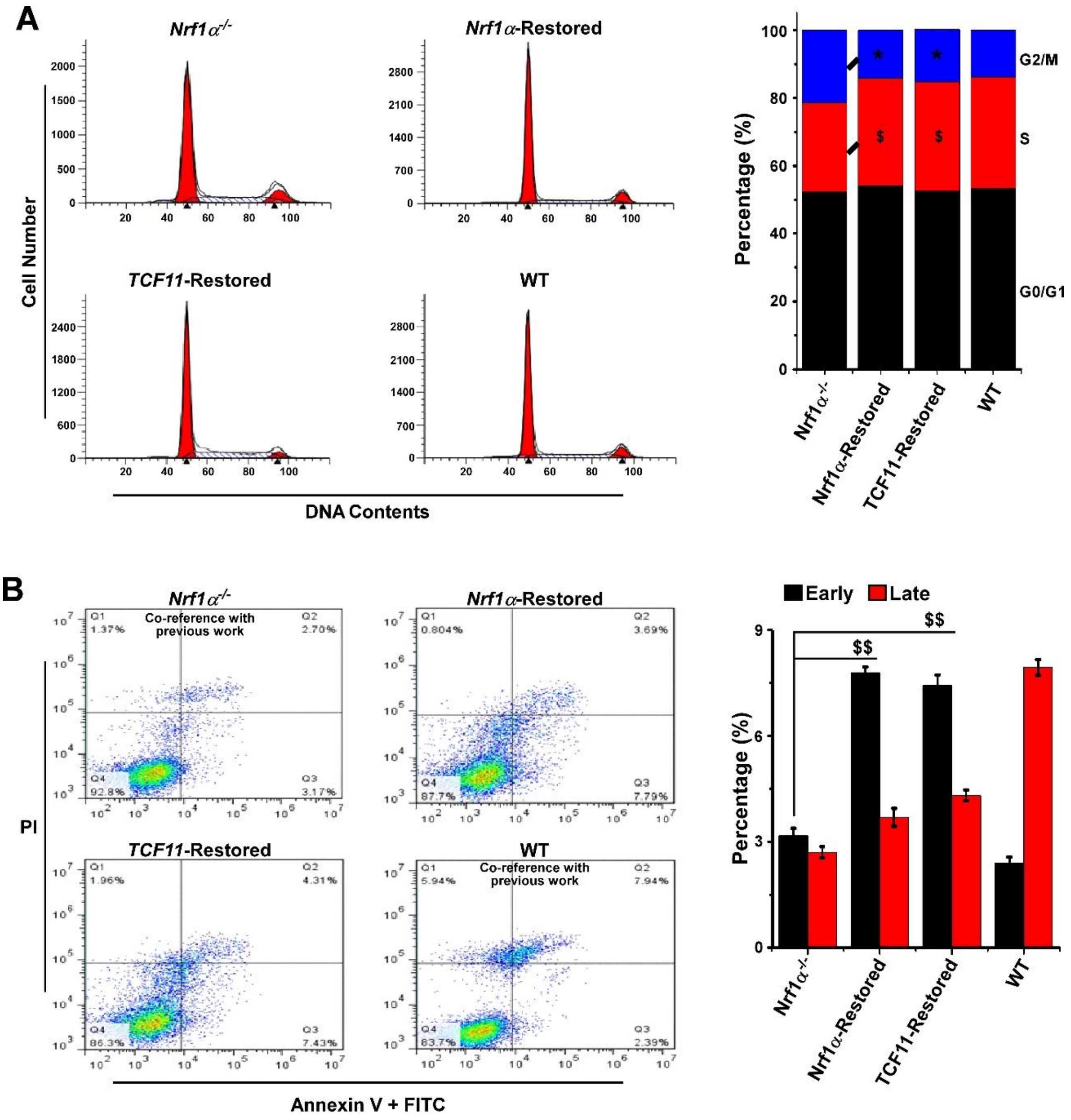


**Figure S7. Alterations in the cell cycle and apoptosis caused by restoration of *Nrf1α* and *TCF11*.** The cell cycle (A) and apoptosis (B) of wild type HepG2 (WT), *Nrf1α*$^{-/-}$, *Nrf1α*-Restored and *TCF11*-Restored cells were examined by using fluorescence-activated cell sorting (FACS). Subsequently, cell distributions were monitored with the BD Accuri C6 software and analyzed by the FlowJo software, and the results were calculated as a percentage (%) of cells examined in different cell-cycle phases or apoptosis status. The data are shown as mean ± SD (n = 9, * $p < 0.05$; $, $p < 0.05$; $$, $p < 0.01$) relative to the corresponding control values obtained form *Nrf1α*$^{-/-}$-derived hepatoma cells.

Figure S8:

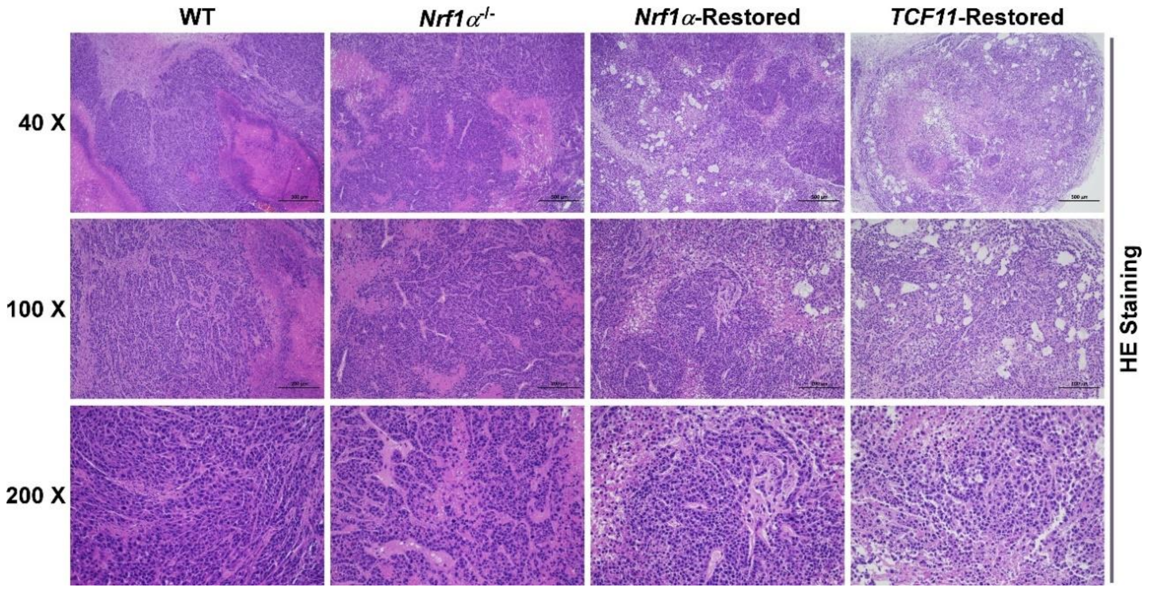


**Figure S8. The HE staining results of tumors in each group.** The histological photographs of indicated tumors were achieved by HE (hematoxylin and eosin) staining.

Figure S9:

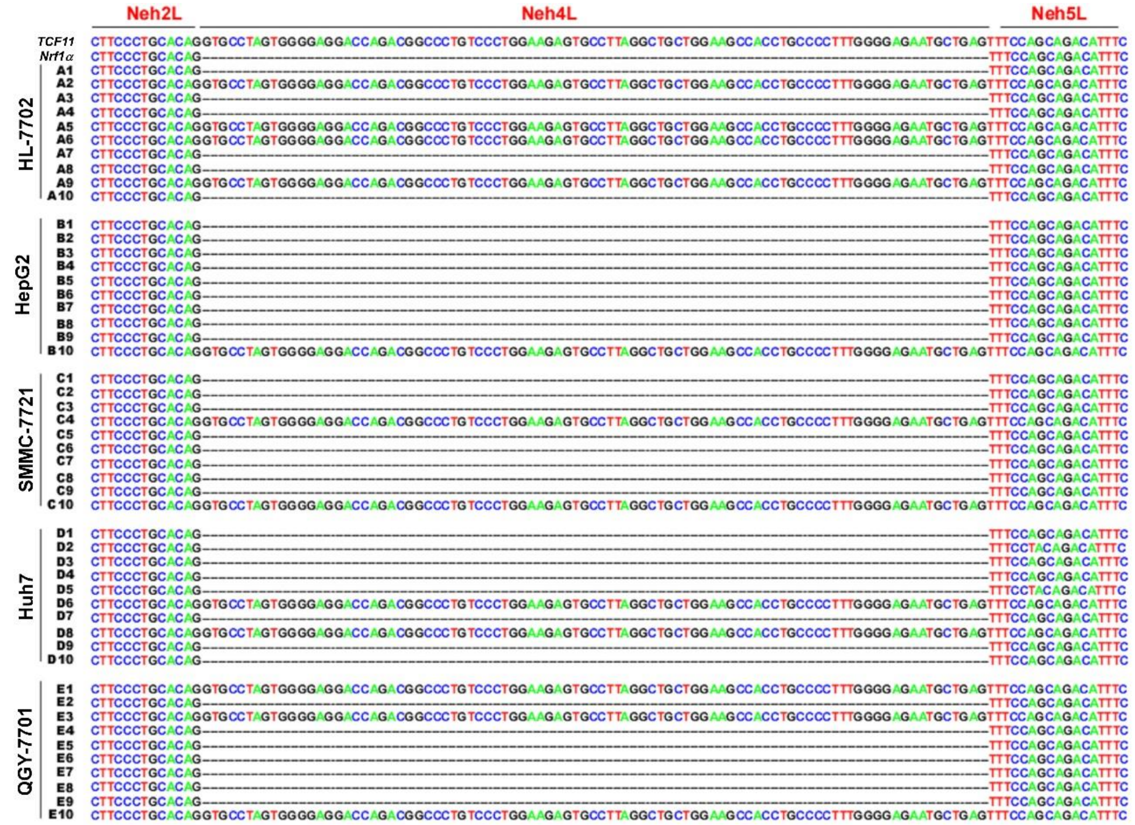


**Figure S9. The relative proportion of *Nrf1α* and *TCF11* in different live cancer cells and normal cells.** Sequencing and blasting result of Nrf1 CDS cloned from different liver cancer cell lines (HepG2, SMMC-7721, Huh7 and QGY-7701) and normal liver cell (HL-7702).
