## Supplemental Tables for "TCF11 Has a Potent Tumor-Repressing Effect than Its Prototypic Nrf1α by Definition of both Similar yet Different Regulatory Profiles, with a Striking Disparity from Nrf2": Supplementary Table S1-Primers.docx

| Oligonucleotides for Construction | | |
| --- | --- | --- |
| Primer Name | Forward (5’-3’) | Reverse (5’-3’) |
| *TCF11* | CGGGGTACCATGCTTTCTCTGAAGAAATACTTAACGGAAG | CCGGATATCCCTTTCTCCGGTCCTTTGGCTTCCTCTCCTG |
| *TCF11^ΔN^* | CGGGGTACCATGGGGGCTGTAGCCCCCCCAGTCAGTGG | CCGGATATCCCTTTCTCCGGTCCTTTGGCTTCCTCTCCTG |
| *Nrf1α* | CGGGGTACCATGCTTTCTCTGAAGAAATACTTAACGGAAG | CCGGATATCCCTTTCTCCGGTCCTTTGGCTTCCTCTCCTG |
| *Nrf2* | CGGGGTACCATGGACTTGGAGCTGCCGCCGCCGGGACTC | CCGGATATCCGTTTTTCTTAACATCTGGCTTCTTACTTTTG |
| Oligonucleotides for qPCR | | |
| Primer Name | Forward (5’-3’) | Reverse (5’-3’) |
| *Nrf1α* | GAGAGCTTCCCTGCACAGTTTC | GACATGAGATCTTGCCACTGCTG |
| *TCF11/*  *TCF11^ΔN^* | CCTTTGGGGAGAATGCTGAGTTTC | GACATGAGATCTTGCCACTGCTG |
| *Nrf2* | AGCTCTCCATATCCCATTCCCTG | GCAGCCACTTTATTCTTACCCCTC |
| *ARFGAP3* | TTACTTTGACGAGCCAGTGGAG | GCCTGTGGTTTTCAGAACTGTTT |
| *ATF2* | TGCTGCTTACAAGTTCTGACTCA | AATTGGCCTGTTAGAGGATGGTG |
| *ATM* | TTTAAGCGCCTGATTCGAGATCCT | CCTGCATCTTTTTCTGCCTGGAG |
| *BARD1* | GCGCTTCTGCACACAGTATATCA | AGCAACTCAAAGGACATCACACA |
| *BLM* | CCCGTATCTTCCCACTACTTTGC | TATCTTTCTACATGTGGCAGACCC |
| *BPTF* | CCAGTAGACCCTAATGATGCACC | TATCTGCCACAAATTCCGTCAGC |
| *BRCA1* | CGGGAGGAAAATGGGTAGTTAGC | TTCTGTCCTGGGATTCTCTTGC |
| *CASP3* | AGAAGATCACAGCAAAAGGAGCAG | AGACTTCTACAACGATCCCCTCT |
| *CAV1* | TTCTCTCTTTCCTGCACATCTGG | CCAACAGCTTCAAAGAGTGGGTC |
| *CCNG1* | TCACTCCTTCAAGAGAACTTGCC | CAATGACATGCCTTCAGTTGAGC |
| *CGREF1* | TCTCCTCTACCTCTTTGCCCTC | GGGTTGGTGGTAGGAGAGTTGG |
| *COL4A6* | TCCTTTCATCGAATGCAGTGGTG | CAGCTTTCAGCGTTTCAGACACA |
| *COL6A2* | GATCGACCAGGACACCATCAAC | ATTTCCAGGCAGCTCACCTTGTA |
| *DICER1* | TACACATGCCTCCTACCACTACA | GTGCTTGGTTATGAGGTAGTCCA |
| *FGF21* | GCCTTGAAGCCGGGAGTTATTC | TATCCGTCCTCAAGAAGCAGCTC |
| *FOSB* | TGACGGCTTCTCTCTTTACACAC | GCAGGTGAGGACAAACGAAGAA |
| *FOXO1* | TGCCCCAGATGCCTATACAAAC | CAGCAACTCCTTCAAGAGTCCAG |
| *GLI1* | GAACCTTCCTACCAGAGTCCCAA | GCCAGGTCCATATGTGTTCACT |
| *GNL2* | CGAAAAAAGTTGGTGTGCGCTAC | TTTTGCGTTTGTGTTTCTGTCCC |
| *HIF1A* | GATACTCAAAGTCGGACAGCCTC | AAAGCGACAGATAACACGTTAGGG |
| *HSP90AA1* | CTGTCTTCTGGCTTCAGTCTGG | TTCTTCAGTTACAGCAGCACTGG |
| *HSP90B1* | GCTGATCAGAGACATGCTTCGAC | AGGGTCAATGTTCAAACTGAGGC |
| *ICAM5* | CTTCAGCTAAATGCCACCGAGA | AGCGTATAGGACACGAAGCTCT |
| *IMPA1* | AACCCCACATGGATCATTGACC | TCCACACAACTGTACACAACTCC |
| *JUNB* | GCAGCTACTTTTCTGGTCAGGG | GACAATCAGGCGTTCCAGCTC |
| *KRAS* | TCTTGGATATTCTCGACACAGCAG | GCAAATACACAAAGAAAGCCCTCC |
| *MAP2K6* | CAACTCCCAGCAGACAAGTTCTC | TGCCACATCTGTTCCTTTGGATT |
| *NF1* | TGATGCTGTGTATTGTCACTCGG | AATGTAAGACTCGGTGCCATTCG |
| *NFKB2* | GACACCCCCCTATCACAAGATG | CTGTTTGGAATCAGACACGTCCC |
| *NRP1* | AAGATCGGGTACAGCAACAACG | GTTGTTGCCCTCAAAAGACTTCG |
| *PALB2* | GCAGCAATCTTGACTTCTGGAAC | CCAGCAAATGAGAGTCTGTACCC |
| *PNN* | GAGCCAGTCTTGACAGTACATCC | GAACTGCTACTGCTACTGCTTCG |
| *PRKCE* | CCACCCTTCAAACCACGCATTA | TGAATTCCTCCTGGTTGATCTGC |
| *PTEN* | TCTTCATACCAGGACCAGAGGA | TCCTTGTCATTATCTGCACGCTC |
| *RAD50* | TGCACATGCTCTGGTTGAGATAA | TCAGAACGTCCTAAAAGCTCCAC |
| *RB1* | GAAGCAACCCTCCTAAACCACTG | TGCTTTTGCATTCGTGTTCGAGT |
| *SMARCA1* | GACAGTGTGTACGAAATGCTCCC | GAGTGTTACAGCGTCTCTGGAAT |
| *SMC3* | ACTGGAGTTGGAATTAGGGTGTC | GGCTACCAAGGATTTCTGTCCAC |
| *SOS1* | ATTCGAGCCCTTTTCACTCAAGC | GCAGATTCTGGTCGTCTTCGTG |
| *STAG2* | AGGAAAACGGAAAGTGGTTGAGG | TGTGGTGTCTGCATCATAACAGG |
| *TGFBR1* | GTGGGAACAAAAAGGTACATGGC | AACATCGTCGAGCAATTTCCCAG |
| *THRB* | CAAGATAGTTTCCTGCTGGCCT | TATCATCCGCAGATCTGTCACCT |
| *USP16* | TTGGCTCCTTTTTGCACCCTTAA | TGACTATTTGCGGTTCTTGCCTT |
| *WNT5A* | ACAACATCGACTATGGCTACCG | TTGTTGTGCAGGTTCATGAGGA |
| *β-actin* | CATGTACGTTGCTATCCAGGC | CTCCTTAATGTCACGCACGAT |
