## Supplemental Tables for "TCF11 Has a Potent Tumor-Repressing Effect than Its Prototypic Nrf1α by Definition of both Similar yet Different Regulatory Profiles, with a Striking Disparity from Nrf2": Supplementary Table S10-prognostic correlation.docx

**Table S9.** Aanalysis of the significance of those potential HCC-associated proteins in relation to prognosis of liver cancers presented in different databases.

| Gene Symbol | ***P*-values** of correlation with prognosis of liver cancer  in different databases used | | | | Prognostic impact |
| --- | --- | --- | --- | --- | --- |
|  | Kaplan-Meier Plotter | UALCAN | OncoLnc | Human Protein Atlas |  |
| ***CHP2*** | < 0.001 | 0.017 | N/A^1^ | N/A | Need to be further verified (Favorable) |
| ***CPS1*** | < 0.001 | 0.011 | < 0.001 | < 0.001 | Favorable |
| ***FOXO1*** | < 0.001 | 0.027 | 0.0778 | < 0.001 | Favorable |
| ***IRS4*** | < 0.001 | N/A | N/A | N/A | Need to be further verified (Favorable) |
| ***NKX2-8*** | < 0.001 | N/A | N/A | N/A | Need to be further verified (Favorable) |
| ***AKR1B10*** | 0.0018 | 0.091 | 0.0146 | 0.0022 | Need to be further verified (Unfavorable) |
| ***EPO*** | < 0.001 | < 0.001 | 0.00414 | < 0.001 | Unfavorable |
| ***MUTYH*** | < 0.001 | < 0.001 | 0.00648 | < 0.001 | Unfavorable |
| ***PKM*** | < 0.001 | N/A | 0.00149 | < 0.001 | Unfavorable |
| ***GP73/GOLM1*** | 0.017 | 0.010 | 0.0554 | < 0.001 | Unfavorable |
| ***GPC3*** | 0.170 | 0.420 | 0.466 | 0.061 | non-significant |
| ***Nrf1*/*NFE2L1*** | 0.620 | 0.220 | 0.487 | 0.052 | non-significant |
| ***Nrf2*/*NFE2L2*** | 0.800 | 0.440 | 0.929 | 0.053 | non-significant |

^1^ No value was found due to insufficient data or low expression levels in liver cancer samples.
